# Myristoylation licenses disordered viral VP4 protein to anchor to and perforate the membrane through phase separation

**DOI:** 10.64898/2026.03.02.708992

**Authors:** Sichao Huang, Fengzhen Deng, Te Liu, Wenjian Li, Peiying Wang, Jiahuan Song, Jingjing Huang, Shiyu Zhang, Jiaxin Liu, Yan Wang, Manjie Zhang, Bin Sun

## Abstract

The VP4 protein of enteroviruses, such as Coxsackievirus B3, is a small, intrinsically disordered protein essential for perforating the host cell membrane during viral entry. A key feature of VP4 is its N-terminal myristoylation, which is required for infectivity in some enteroviruses but dispensable in others, suggesting a complex and context-dependent role that is not fully understood. The precise biophysical mechanisms by which this lipid anchor enables a disordered protein to breach a membrane remain unresolved. Here, using Coxsackievirus B3 VP4 as a model system and integrating multi-scale molecular dynamics simulations with confocal microscopy, we demonstrate that myristoylation is not a simple membrane tether but a multi-functional regulator that orchestrates VP4 activity through distinct, hierarchical roles. First, it provides the necessary hydrophobic anchor to recruit the disordered VP4 to the membrane interface. Second, the myristoyl group acts as a key molecular driver that promotes the liquid-liquid phase separation of VP4, leading to the formation of dynamic condensates on the membrane surface. These condensates actively remodel the membrane, generating substantial curvature that, in turn, lowers the free energy barrier for VP4 penetration. Furthermore, we find evidence that the myristoyl group plays a third role in stabilizing the final transmembrane pore. Our findings establish a novel paradigm where a single lipid modification empowers a disordered viral protein to form a functionally potent condensate that mechanically primes and physically breaches the target membrane, a mechanism that may explain the conditional myristoylation requirement across enteroviruses.

## 1 Introduction

Non-enveloped viruses, also called naked viruses, are composed solely of the viral genome and proteins, lacking an external lipid envelope. During infection by these viruses, a vital step is crossing the host cell membrane to deliver their genome into the cell. Unlike enveloped viruses, which breach the host cell membrane via membrane fusion [1], non-enveloped viruses rely on their own capsid proteins to permeate the membrane. Picornaviruses, a major family of nonenveloped pathogens that includes coxsackievirus B3 (CVB3), rhinoviruses, and poliovirus, cause various diseases ranging from the common cold to life-threatening inflammation [2, 3]. They are formed capsids from 22 to 30 nm in diameter and contain a single-stranded RNA genome of approximately 7500 nucleotides [4]. The picornavirus capsid is composed of structural proteins VP1, VP2, VP3, and VP4, organized with pseudo-T=3 icosahedral symmetry. During infection, a cascade of structural changes in the viral capsid, referred to as uncoating, occurs. This is followed by the release of VP4 from the capsid. Then, the released VP4 participates in forming a size-selective pore in the host membrane, enabling genome translocation [5, 6]. To date, the indispensable role of the VP4 protein for infectivity across many picornaviruses is well-documented [7, 8].

Despite the conserved functional role of VP4 in membrane breaching across the picornavirus family, these proteins exhibit two distinct functional paradigms regarding their dependence on N-terminal myristoyl modification. Myristoylation is a lipid post-translational modification with the well-established function of anchoring proteins to membranes [9]. For some picornaviruses, myristoylation is a prerequisite for VP4 function, while for others, it is dispensable. For example, in enteroviruses such as CVB3 and Enterovirus 71 (EV71), infectivity is strictly dependent on the myristoylated form of VP4 for membrane penetration [10, 11]. Loss of VP4 myristoylation through mutation or inhibitor treatment significantly impairs membrane breaching [10]. In contrast, for viruses such as Hepatitis A virus (HAV), Aichi virus (AiV), Minute virus of mice (MVM), and Triatoma virus (TRV), VP4 functions effectively without this modification [12–15]. The molecular basis for this differential myristoylation-dependence of VP4 function remains elusive, particularly for viruses that require it. Understanding this is of significant therapeutic interest, as the indispensable role of myristoylated VP4 for infectivity presents a potential drug target. For instance, antibodies targeting the myristoylated N-terminus of Human Rhinovirus (HRV) VP4 demonstrate significant cross-serotype neutralization [16].

Accumulated evidence suggests that VP4’s dependence on myristoylation is likely correlated with its structural state. Structurally, myristoylation-dependent VP4 proteins are predominantly intrinsically disordered, with large segments unresolved in capsid structures [17, 18]. For example, X-ray crystallography shows that enterovirus 71 (EV71) VP4 is disordered within the viral particle [17]. Similarly, in cryo-EM structures of CVB, HRV, and enterovirus D68 (EVD68) capsids, the N-terminal region of VP4s are partially resolved, and the remaining domain almost exclusively adopt loop configurations [18–20]. In contrast, VP4 proteins that function effectively without myristoylation tend to adopt stable, pre-folded helical structures [12, 13]. From an amino acid composition perspective, non-myristoylated VP4 peptides appear to be more hydrophobic than their myristoylated counterparts [21]. It has been speculated that the hydrophobic myristoyl moiety likely spearheads membrane integration, thereby reducing the requirement for extensive hydrophobic interactions from the rest of the VP4 protein [21]. This may suggest that myristoylation is essential to compensate for the lack of inherent structure and hydrophobicity in disordered VP4 proteins. However, the precise molecular mechanism by which myristoylation enables a disordered protein like VP4 to efficiently breach membranes remains elusive.

In this study, we use the CVB3 VP4 protein as a model system. By combining multiscale molecular dynamics simulations with experimental assays, we provide a molecular picture of how VP4’s structural flexibility and myristoyl modification interact to enable its function. First, we confirm the intrinsically disordered nature of CVB3 VP4 and elucidate how this flexibility works in concert with the myristoyl group to enable membrane interaction. We then demonstrate that myristoylation further drives the disordered VP4 to form biomolecular condensates on the membrane surface. These condensates, in turn, facilitate the insertion of individual VP4 proteins into the lipid bilayer by lowering the energy barrier. Finally, we explore the functional endpoint of this process, providing evidence that the N-terminus of VP4 adopts a helical conformation upon membrane insertion and that myristoylation contributes to stabilizing putative multimeric pore structures. Our findings revealed the mechanism wherein myristoylation licenses a disordered viral protein for membrane penetration not merely as a simple anchor, but as a key driver of functional, collective behavior through phase separation. This work provides a unifying structural and biophysical framework to explain the mystery of myristoylation-dependence in viral membrane-penetrating proteins.

## 2 Methods

### 2.1 Sequence retrieval and structural preparation of VP4 protein

The full-length sequence of CVB3 VP4 was obtained from the NCBI database (GenBank ID: AAA42931), which is consistent with the VP4 plasmid preserved in our laboratory. The complete three-dimensional structure of VP4 was predicted using the Swiss-Model server [22] with the crystal structure PDB 4GB3 as template [23]. The target template sequence identity was determined to be 89.58%, with a Global Model Quality Estimation (GMQE) score of approximately 0.7. pKa calculations were performed using the PropKa server [24], which predicted that the histidine residues at positions 13 and 26 are prone to protonation at neutral pH (pH 7.0). Accordingly, to align with the default neutral pH conditions of the molecular dynamics (MD) simulations, these two histidine residues were modeled in their protonated state (HSE). Finally, the VP4 structure was submitted to the CHARMM-GUI web server, developed by the Wonpil group[25], to generate input files for MD simulaitons.

### 2.2 Atomistic molecular dynamics simulations of isolated VP4

The CHARMM-GUI server was used to generate the input files of atomistic conventional MD simulations [25]. Proteins were described by the CHARMM36m force field [26] and solvated in a TIP3P water box with a 12 Å buffer distance, resulting in a box of the size 98.852 Å × 98.852 Å × 98.852 Å. The NaCl ions were added to the system to maintain a salt concentration of 0.15 M. For the myristoylated VP4 structure, the force field parameters for the myristoylated Glycine were adapted from the as-prepared parameters from CHARMM-GUI. The system first undergo 50,000 steps of energy minimization, the initial 200 steps used the steepest descent algorithm, followed by the conjugate gradient algorithm for the remaining steps. The minimized system was then heated to 300 K through a two-step procedure: heating from 0 to 100 K over 0.1 ns in the NVT ensemble, followed by heating from 100 to 300 K over 0.5 ns in the NPT ensemble. During the heating process, harmonic constraints with a force constant of 5 kcal/mol/Å^2^ were applied to the backbone atoms. After heating, the system underwent 1 ns of equilibrium simulation in the NPT ensemble at 300 K, with the force constant reduced to 1 kcal/mol/Å^2^. Finally, three independent 1 *µ*s production runs were performed. During the simulations, all bonds involving hydrogen were constrained using the SHAKE algorithm [27], temperature was controlled by the Langevin thermostat, and long-range electrostatic interactions were handled using the Particle Mesh Ewald (PME) method. The time step was set to 2 fs, with a non-bonded cutoff of 10 Å. The Amber20 package was used to the above atomistic conventional simulations [28].

### 2.3 Martini Coarse-grained MD (CGMD) simulations of VP4-membrane interaction and VP4 condensation

The Martini 3.0 coarse-grained (CG) model was employed to simulate the protein-membrane interaction [29]. The coarse-grained structures of VP4 were generated from the corresponding atomistic structures sampled from atomistic MD simulation using the Martinize2.py script as described on the MARTINI website. The topology of myristoylated Glycine was generated using the CG Builder program, and the Martini force field parameters was obtained from previous research [30]. Since the VP4 protein is intrinsically disordered, we did not include any elastic networks on the protein, and we termed this mode as VP4-IDP model. We additionally built two VP4 CG models that are partially rigid, and fully rigid, respectively. For the VP4-partial model (which has 132 elastic bonds), the bead pairs involved in the elastic bonds were determined using the full-length VP4 structure obtained via Swiss homology modeling based on the partially resolved VP4 crystal structure in PDB 4GB3. The default Martini elastic bond criteria, with lower and upper elastic bond cutoffs of 0.5 and 0.9 nm, respectively, were used, resulting in 132 elastic bonds. For the fully rigid VP4 model, the fully relaxed VP4 structure from the atomistic MD simulation of VP4 was used to determine the elastic bonds, resulting in a total of 206 bonds using the same cutoffs. All elastic bonds have a force constant of 2.39 kcal/mol/Å^2^ with a cutoff distance of 11 Å. The membrane is composed of POPC CG molecules, and has a dimension of ∼200 Å × 200 Å in the *xy* direction. Since long-range electrostatic interactions between proteins become centrosymmetric around 40 Å [31], the VP4 protein(s) was randomly positioned at approximately 40 Å above the lipid bilayer surface. At such distances, electrostatic interactions between proteins are weak and centrosymmetric [32, 33], which could ensure that the electrostatic interactions between VP4 proteins (including the MYR) and membrane lipids are unidirectional. Thus, while the native driving force for VP4 aggregation is preserved, the recruitment of these proteins to the membrane is initially governed by thermally driven motions, avoiding any bias that artificially enhance the membrane-recruitment effect while obscure the aggregation-promoting role of the MYR moiety. Martini water molecules and NaCl ions correspond to 0.15 M ionic strength were added into the system.

The system first underwent energy minimization for 2000 steps using the steepest descent algorithm, followed by heating to 300 K in the NPT ensemble over 1 ns with harmonic constraints of 2.39 kcal/mol/Å^2^ applied to the protein. The equilibrated system then underwent 10 *µ*s production MD simulation in the NPT ensemble at 300 K using a 20 fs time step. Temperature was controlled using the velocity rescale (V-rescale) thermostat, and pressure was maintained with the Parrinello-Rahman barostat. For each system, three independent 10 *µ*s CG MD replicas were conducted using Gromacs 2020.4 [34]. The back mapping from CG model to the all-atom model was performed according to the methods described in Martini website [35]. The size distribution of VP4 protein clusters was calculated using the *gmx clustsize* command, with a maximum distance cutoff of 5.5 Å for cluster identification [36].

The mean curvature of the membrane was calculated by using the MDAnalysis tools [37]. The phosphate bead groups (PO4 beads) in the upper leaflet were selected as reference points to reconstruct the membrane surface and compute its corresponding mean curvature. The analysis grid dimensions were defined by the size of the periodic simulation box in the *xy* plane (∼200 Å × 200 Å). To balance computational efficiency and avoid overfitting, this planar grid was uniformly discretized into 10 × 10 bins, with each bin side length being ∼20 Å. Periodic boundary wrapping was enabled to ensure all atoms were correctly positioned within the primary unit cell prior to surface fitting. For each frame in the trajectory, the mean curvature field was computed from the reconstructed surface using standard differential geometry methods. The final membrane curvature map was obtained by temporally averaging the curvature values across all frames in the analyzed ensemble.

### 2.4 One-dimensional coarse-grained umbrella sampling to study the translocation of individual VP4 proteins from the condensate into the membrane

Umbrella sampling (US) simulations were conducted to estimate the potential of mean force (PMF) for the insertion of an individual VP4 protein from the condensate into a POPC membrane. These simulations were performed at the Martini coarse-grained (CG) level, using the final frame from a conventional Martini CG MD trajectory as the starting structure. In this way, the membranes in the PMF calculations retain their curvature, as reshaped by the monomer, hexamer, and 20-mer VP4 proteins. The VP4 molecule selected for membrane translocation was one with its myristoyl moiety already inserted into the membrane. The reaction coordinate (RC) was defined as the distance between the center of mass (COM) of the N-terminal myristoylated glycine (Myr-Gly) and the COM of the lipids in the lower membrane leaflet. This RC definition yields positive values when the myristoyl moiety is distal to the lower leaflet, a value near zero when it approaches the leaflet plane, and negative values when the moiety penetrates the lower leaflet, indicating substantial immersion of the VP4 protein into the membrane bilayer. The RC was explored from approximately +20 Å to −30 Å, divided into windows of 1 Å width. A harmonic restraint with a force constant of 10 kcal/mol/Å^2^ was applied in each window. The initial structure for each window was taken from the final frame of the preceding window. Production molecular dynamics (MD) simulations were run for 100 ns per window in the NPT ensemble at 300 K. The last frame of the preceding window served as both the starting structure and the reference configuration for the next window. During the PMF calculations, except for the pulling force acting on the selected VP4 protein, no other forces are applied to the system. The resulting PMF profiles were computed using the WHAM program [38]. Simulations were preformed using Gromacs 2020.4 software [34].

### 2.5 Well-Tempered metadynamics (WT-MetaD) simulations studying the helix-folding of the N-terminal VP4

All-atom WT-MetaD simulations were performed using the Gromacs-2022.5 package patched with the PLUMED 2.8 plugin [39] to calculate the folding free energy of the N-terminal 20 residues of the VP4 protein, with and without myristoylation modification, respectively. The reaction coordinate (RC) was defined as the residual helicity of this VP4 N-terminal region, representing the fraction of residues forming an ideal *α*-helix; this value ranges from 0 (fully unwound) to 1 (completely *α*-helical). We studied folding both in aqueous solution and within a POPC membrane bilayer.

The initial configuration of the N-terminal fragment, extracted as a coil from the last frame of atomistic simulated full-length VP4 structure, was used for both scenarios. For the aqueous simulation, the fragment was solvated in a TIP3P water box (size: ∼60 Å × 60 Å × 60 Å) containing 0.15 M KCl. For the membrane environment simulation, the fragment was embedded at the center of a POPC bilayer (size: ∼90 Å × 90 Å in the *xy*-plane), with TIP3P water layers containing 0.15 M KCl (thickness: ∼30 Å) added above and below the membrane. Specifically, initial VP4 fragment was placed at the interface between the upper and lower lipid leaflets (i.e., approximately at the center of the membrane, see Fig. S9). We verified that during WT-MetaD, the VP4 fragment (including the MYR moiety) remained within the lipids. Since our purpose is to explore whether the VP4 fragment has a different helix-folding propensity in water versus in a hydrophobic environment, such a WT-MetaD setup, in which the VP4 fragment is completely embedded within the lipids, fulfills our purpose. All systems were described using the CHARMM36m force field [26]. Minimization, heating, and equilibration followed the aforementioned MD protocol. WT-MetaD was initiated from the equilibrated system. The bias potential had an initial height of 0.96 kcal/mol, a Gaussian width (*σ*) of 0.1 (in RC units), and was deposited every 0.5 ps. A bias factor of 20 was used, and the temperature was maintained at 300 K. For each system, three independent 0.5 *µ*s WT-MetaD production runs were performed.

### 2.6 Two-dimensional all-atom umbrella sampling to explore the correlation between helical content with embedding depth of the N-terminal fragment of VP4 within membrane

The highest helical content configuration of the N-terminal 20 residues of VP4 sampled via WT-MetaD simulations in membrane was used as starting structure to initiate a 500 ns of production MD. The last frame of such MD trajectory serves as the starting structure for the two-dimensional umbrella sampling simulations. The first RC was defined as residual helical content of the fragment, which varies from 0 to 1 and was binned into 0.1-width windows. The second RC was defined as the distance between the center of mass (COM) of VP4 and the COM of the headgroup region of the lower leaflet, ranging from 38.6 Å to 0 Å and was binned into 0.6 Å width. Notably, this distance is not the minimal distance between VP4 atoms and the membrane surface. Harmonic restraints applied on first RC was 4.18 kcal/mol/Å^2^, and was 2.39 kcal/mol/Å^2^ for the second RC. In total, 726 sampling windows were sampling in the 2D RC space. For each window, a 5 ns production simulation was carried out in the NPT ensemble at 300 K, resulting in an aggregate sampling time of 3.63 *µ*s. The 2D PMF profiles were computed using the WHAM program. Simulations were preformed using Gromacs 2020.4 software [34].

### 2.7 All-atom MD simulations to investigate the stability of the VP4 pore within a POPC membrane

Since VP4 has been reported to form a six-fold symmetric pore within the membrane, we manually constructed model pores using helices formed by the N-terminal 20 residues of VP4 (both with and without myristoylation). The helical conformation for these fragments was selected from configurations sampled via WT-MetaD, characterized by a helical content of approximately 0.8. Six such helices were vertically immersed in a POPC membrane measuring ∼100 Å × 100 Å in the *xy* plane, with the inter-helical distance adjusted to allow only van der Waals contacts. The pore system was solvated with TIP3P water and NaCl, with water layers added above and below the membrane. Conventional MD simulations using the CHARMM36m force field were then performed via Gromacs to assess the stability of pores formed by myristoylated and non-myristoylated VP4 N-terminal fragments. Following system equilibration, three independent 0.5 *µ*s production MD runs were conducted for each case. A summary of all molecular dynamics simulations performed in this study is provided in Table S1.

### 2.8 Cells, main regeagents and plasmids

HEK-293T cells were maintained in modified Eagles medium supplemented with 10% fetal bovine serum at 37 řC in 5% CO2. Escherichia coli BL21 was purchased from TransGen Biotech (Beijing,China). Lipofectamine 2000 transfection reagent was purchased from GeneCopoeia (Rockville, MD, USA). 1,6-Hexanediol was purchased from Med Chem Express (Monmouth Junction, NJ). VP4-WT-GFP, VP4-G2A-GFP and PET28a-VP4-WT plasmids were all preserved in our lab at -80řC.

### 2.9 Circular Dichroism

The VP4 protein was expressed and purified for structural analysis. Briefly, the pET28-His-VP4 plasmid was transformed into E. coli BL21(DE3) cells and induced with 0.5 mM isopropyl *β*-D-1-thiogalactopyranoside (IPTG; Clontech, CA, US). After expression in Escherichia coli, the recombinant VP4 proteins were purified via Ni-affinity chromatography using a His-Trap column, and the purified fractions were dialyzed into assay buffer (50 mM Tris pH 8.0, 200 mM NaCl). Subsequently, circular dichroism (CD) spectra of the purified VP4 were then acquired on a J-815 spectropolarimeter (Jasco) over the wavelength range of 190260 nm. The protein sample was prepared at a concentration of 50 *µ*M in 10 mM sodium phosphate buffer.

### 2.10 Live-cell imaging

HEK-293T cells grown in glass-bottomed cell culture dishes for 24h before transfection. At about 70% confluence, cells were transfected with EGFP-VP4-WT or EGFP-VP4-G2A using Lipofectamine 2000, respectively. At 12 h after transfection, the 4’,6-diamidino-2-phe-nylindole (DAPI; Coolaber, Beijing, China; 1:750 dilution) was used to stain the cell nucleus under dark environment and room temperature for 25 min. Finally, the cells was scanned under a confocal laser scanning microscope (Zeiss LSM 800, German). Timelapse images of EGFPVP4 were acquired at 60 s intervals under consistent environmental and instrumental settings.

### 2.11 Analysis of the effect of 1,6-Hexanediol on VP4 phase separation

To test the effect of 1,6-hexanediol on VP4 phase separation in cells, cells were transfected with the GFP-VP4-WT plasmid for 12 hours. Subsequently, 1,6-hexanediol was diluted in the cell culture medium to a final concentration of 10% (w/v). The original medium was gently replaced with the 1,6-hexanediol-containing medium, and and images were recorded at 1 min intervals.

### 2.12 Fluorescence recovery after photobleaching assay

FRAP analysis was performed using the FRAP module of the confocal laser scanning microscope. HEK-293T cells were transfected with the VP4-GFP plasmid for 12 hours prior to imaging. Circular condensates of VP4-GFP were photobleached within regions of interest using a 488 nm laser, reducing fluorescence intensity to 10%20% of initial levels. Fluorescence recovery was monitored by acquiring time-lapse images every 3 seconds for 240 seconds. The fluorescence intensity within the bleached areas was background-corrected and normalized to pre-bleach values.

## 3 Results and Discussion

### 3.1 Disordered nature of the VP4 protein from CVB3

The VP4 protein of Coxsackievirus B3 (CVB3) is a 68-residue polypeptide located inside the viral capsid prior to its externalization during infection (Fig. 1A). In crystallographic structures, VP4 is typically resolved as part of a complex with VP1, VP2, and VP3 proteins, forming a basic capsid subunit (Fig. 1B). Notably, the N-terminal region of VP4 is absent from the crystal structure due to a lack of electron density, while the remaining resolved residues adopt a loop-like conformation. This indicates a high degree of structural flexibility both at the full-length protein level and specifically at the N-terminus, where the myristoylation site is located (Fig. 1B).

**Figure 1:**
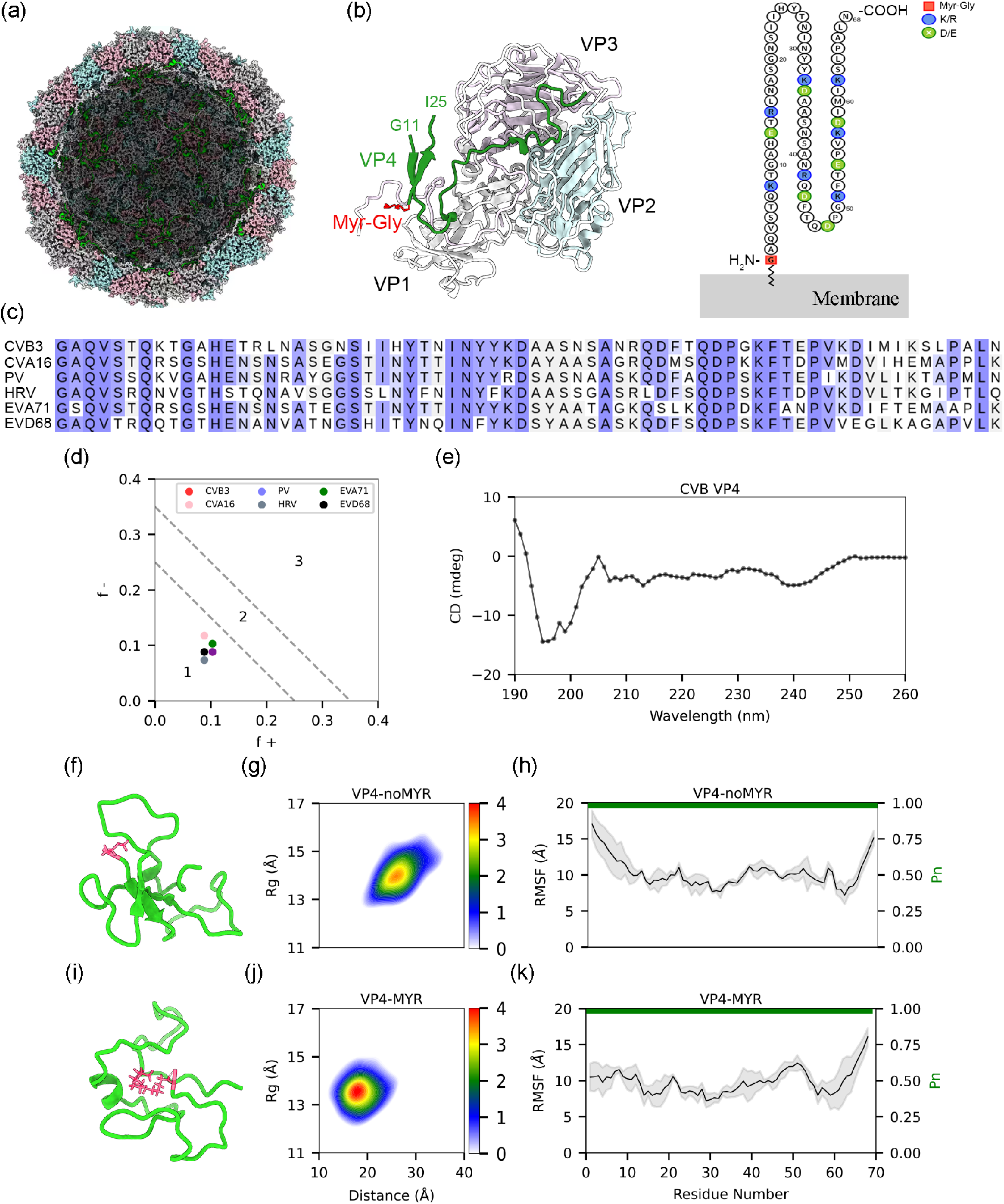
The intrinsically disordered nature of the CVB3 VP4 protein. (A) Cross-sectional view of the CVB3 capsid (rendered from PDB 4GB3 [23]) showing the interior location of the VP4 protein (green). Other structural proteins VP1, VP2, and VP3 are colored gray, cyan, and pink, respectively. (B) Crystal structure of the basic capsid subunit formed by VP1VP4 (PDB 4GB3 [23]). Residues 1224 of VP4 are unresolved. A topology diagram highlights the myristoylation site and a hypothesized working model for myristoylenabled VP4 function. (C) Sequence alignment of VP4 from six viruses that require myristoylation for function. (D) Predicted structural states of the six VP4 proteins plotted on the Das-Pappu diagram (region 1: weak polyampholytes/polyelectrolytes). (E) Circular dichroism spectrum of CVB3 VP4 in phosphate buffer (pH 7.0). (F, I) Representative conformational snapshots of VP4 without (F) and with (I) the myristoyl modification. VP4 residues 268 are colored green; the myristoyl group is red. (G, J) Conformational ensembles projected onto a 2D plane defined by the endtoend distance and radius of gyration (Rg). (H, K) Perresidue RMSF and the total nonhelical secondarystructure propensity (P*n*) derived from the simulations. Shaded areas represent the standard deviation from three independent 1 *µ*s MD replicates.

Sequence analysis reveals that the myristoylation site region is highly conserved among VP4 proteins known to undergo this modification (Fig. 1C). In contrast, VP4 proteins that function independently of myristoylation show lower overall sequence conservation (Fig. S1). To further assess their structural propensity, we calculated the fraction of positive and negative charges for myristoylated VP4 sequences and plotted them on the Das-Pappu diagram [40]. These proteins consistently fall within the region characteristic of weak polyampholytes and polyelectrolytes (region 1), suggesting a natural tendency to adopt compact conformational ensembles such as tadpoles or globules (Fig. 1D). We further confirmed the disordered state of CVB3 VP4 experimentally using circular dichroism (CD) spectroscopy. The spectra are dominated by a strong negative band near 200 nm, characteristic of a random coil conformation, with minimal *α*-helical or *β*-sheet content (Fig. 1E). This finding aligns with the crystallographic evidence and previous reports [19], establishing VP4 as an intrinsically disordered protein (IDP) in solution.

To characterize the conformational ensemble of VP4 and probe the influence of myristoylation, we performed extensive all-atom conventional Molecular Dynamics simulations (3 Œ 1 *µ*s each) of the unmodified (VP4-noMYR) and myristoylated (VP4-MYR) protein. Representative structures show that VP4-noMYR samples more extended conformations, whereas VP4-MYR adopts collapsed structures that wrap around the hydrophobic myristoyl moiety (Fig. 1F, I). Projecting the simulation trajectories onto a two-dimensional space defined by the end-to-end distance and the radius of gyration (Rg) quantifies this difference: the VP4-noMYR ensemble centers around an end-to-end distance of ∼26 Å and an Rg of ∼14 Å, both slightly larger than the corresponding values for VP4-MYR (∼17 Å and ∼13.5 Å, respectively; Fig. 1G, J). This indicates that the myristoyl group induces a moderate compaction of the VP4 conformational ensemble.

Despite this compaction, both forms remain highly disordered. This is evidenced by similarly high backbone root-mean-square fluctuation (RMSF) values (∼12 Å) across the sequence for VP4-noMYR and VP4-MYR (Fig. 1H, K). Furthermore, both variants exhibit a predominantly coil-like secondary structure as the total nonhelical secondarystructure propensity P*_n_* approaches 1.0 in both cases. DSSP algorithm-based [41] secondary structure analysis further confirmed that VP4-MYR and VP4-noMYR essentially did not form helices (Fig. S2). In summary, our combined computational and experimental data conclusively demonstrate that the CVB3 VP4 protein is intrinsically disordered in solution.

### 3.2 Myristoylation facilitates membrane anchoring of VP4, a process that requires high conformational flexibility of the protein

The primary function of protein myristoylation is to promote protein-membrane interaction [42]. For some picornaviruses VP4s, myristoylation has been reported to enable VP4-membrane interaction, but the molecular basis remains elusive. This is particularly intriguing given VP4’s highly dynamic nature, and the observation that its myristoyl moiety is relatively buried within the protein in solution from CVB3 (Fig. 1F). To investigate the role of myristoylation in VP4-membrane interaction, we placed a monomeric VP4 protein approximately 40 Å away from the membrane surface and conducted coarse-grained MD simulations to monitor the association process (Fig. 2A). The time-dependent protein positions relative to membrane shows that myristoylated VP4 quickly binds and stabilizes on the membrane within 0.72 *µ*s, while the non-myristoylated counterpart merely bounces upon contact (Fig. 2B). In the membrane-anchored state, the N-terminal myristoyl moiety is deeply inserted (∼10 Å) into the membrane, while the protein’s center of mass (COM) lies on the membrane surface (Fig. 2C) steadily. In contrast, VP4-noMYR approaches the membrane more slowly and only forms transient contacts before dissociating; the Glycine residue (the myristoylation site) and the protein’s COM remain consistently *>*10 Å from the membrane, highlighting the inability of non-myristoylated VP4 to anchor to the membrane. Atomistic structure backmapped from membrane-bound VP4-MYR state revealed extensive contacts between the N-terminal myristoyl moiety and the membrane lipid tails, while very few contacts were formed between VP4-noMYR and lipids (Fig. S3). These data suggest that myristoylation plays a pivotal role in facilitating VP4 membrane binding by inserting directly into the lipid bilayer, providing a molecular mechanism for the previously reported role of myristoylation [10, 43].

**Figure 2:**
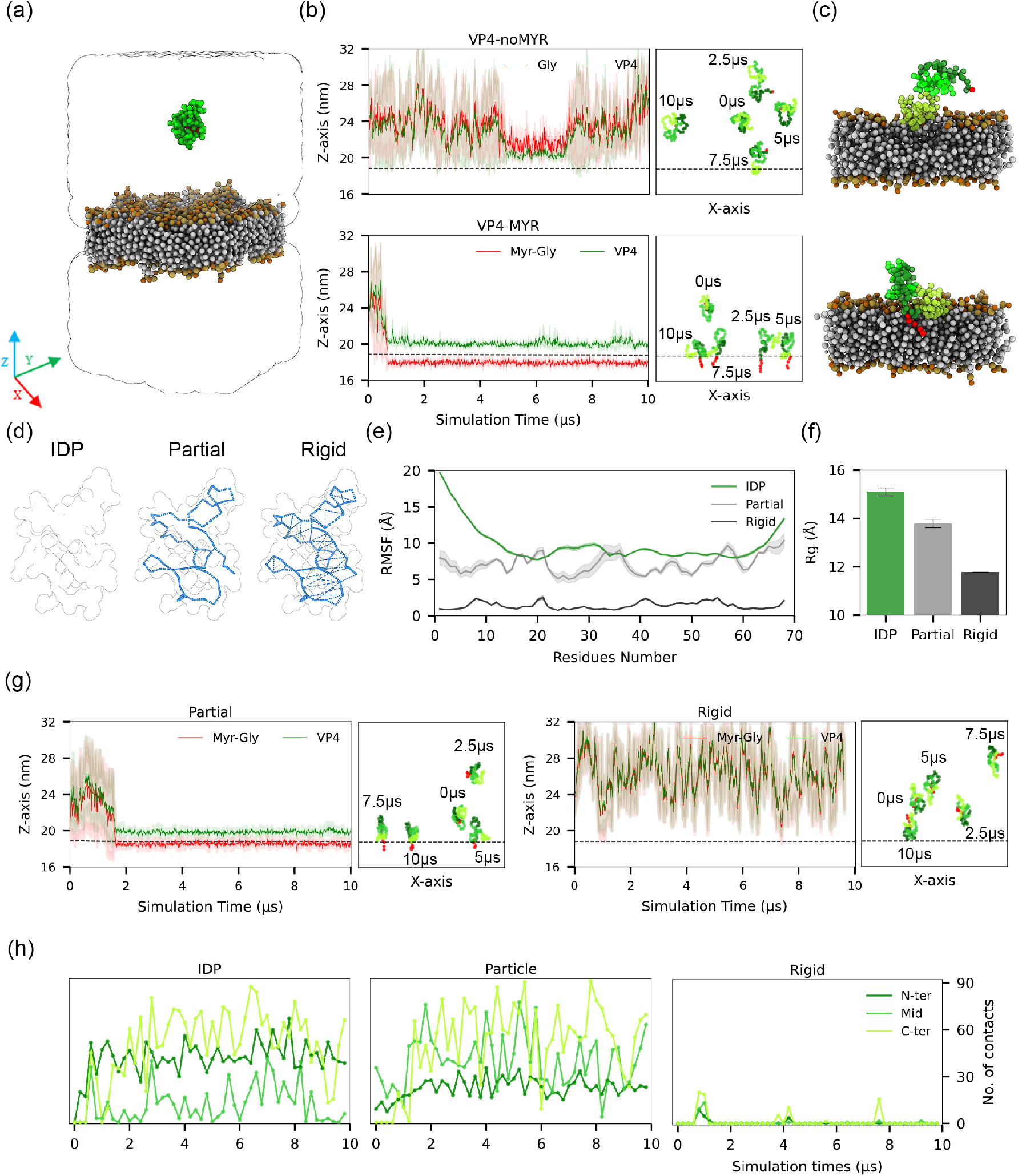
MD simulations of monomric VP4 binding to membrane. (A) Schematic of CGMD simulation setup. VP4 protein is colored green with myristoylation-modificated Glycine colored red. (B) Time-dependent z-coordinates of the COM for the myristoyl group and the VP4. Shaded area represents the standard deviation from 3*×*10 *µ*s CGMD simulations for each condition. The dashed line indicates the average z-coordinate of the COM for the polar headgroups of POPC lipids in the upper leaflet. Snapshots taken at 2.5 *µ*s intervals from a representative 10 *µ*s trajectory are projected onto the XZ plane to illustrate the binding process. (C) Representative structures of VP4 in its membrane-bound state, showing minimal separation from the membrane. (D) Schematic illustrating the construction of two VP4 models with reduced conformational flexibility: VP4-partial and VP4-rigid. These were generated by introducing increasing amounts of intramolecular elastic restraints (blue lines). (E, F) RMSF and Rg of the VP4 models from simulations in solution, confirming their reduced conformational flexibility to the native VP4 IDP. (G) Time-dependent z-coordinates of the VP4-partial and VP4-rigid models during membrane interaction. Snapshots from the simulations are projected onto the XZ plane. (H) Time-dependent number of contacts between the membrane and the N-terminal (residues 1-20, deep green), middle (residues 21-40, green), and C-terminal (residues 41-68, light green)1s3egments of VP4.

Given that VP4 is an IDP, we next explored whether the membrane-anchoring function of the myristoyl group depends on VP4’s structural flexibility. We built two models with reduced flexibility: one with partially reduced flexibility (“VP4-partial”) and one that was essentially rigid (“VP4-rigid”)(Fig. 2D; see Methods for details). The RMSF and Rg of the isolated proteins confirmed that these models had the intended reductions in conformational flexibility compared to native VP4 (Fig. 2E-F). We then simulated the membrane binding of these two artificial models in the presence of the myristoyl group. The VP4-rigid model completely lost its ability to bind the membrane (Fig. 2G). The VP4-partial model retained some binding ability, but it bound much more slowly (first binding at ∼1.8 *µ*s vs. ∼0.72 *µ*s) and more shallowly (myristoyl insertion depth of ∼2.5 Å vs. ∼10 Å) than native VP4.

To understand how reduced flexibility compromises membrane anchoring, we divided VP4 into N-terminal, middle, and C-terminal segments and calculated the time-dependent contact number for each with the membrane. Overall, native VP4 formed the highest number of total contacts (Fig. 2H). As flexibility decreased, the VP4-partial model formed moderate contacts, and the VP4-rigid model formed almost none (Fig. 2H). The reduction in contacts was primarily contributed by the N-terminal segment, suggesting that the high conformational flexibility of VP4 is a prerequisite for the myristoyl group to effectively anchor the protein to the membrane. Notably, for native VP4, the myristoylated glycine (Myr-Gly) contributed more than half of the N-terminus’s contacts with the membrane (Fig. S4), demonstrating the critical role of the myristoyl group. Meanwhile, the C-terminus of myristoylated native VP4 also contributed a significant number of contacts, indicating that this region, which is rich in hydrophobic residues, plays an auxiliary role in membrane interaction. This is consistent with previous reports [9, 44, 45].

### 3.3 Myristoylation promotes VP4 condensate formation on the membrane surface

To verify the membrane-anchoring ability of the myristoyl group in a cellular environment, we expressed EGFP-tagged wild-type (WT) VP4, which is myristoylated, and a G2A mutant VP4, which lacks the myristoylation site, in HEK-293T cells. As shown in Fig. 3A, WT VP4 is enriched at the membrane, while the G2A mutant is diffuse in the cytoplasm, confirming the critical role of myristoylation in localizing VP4 to the membrane. Furthermore, WT VP4 was not evenly distributed on the membrane but instead aggregated to form discrete puncta, suggesting that myristoylated VP4 undergoes liquid-liquid phase separation (LLPS). To confirm the LLPS phenomenon, we performed time-lapse imaging of WT EGFP-VP4 and observed the progressive fusion of VP4 puncta and their dynamic movement over time (Fig. 3B). A fluorescence recovery after photobleaching (FRAP) assay demonstrated rapid fluorescence recovery within the condensates, confirming their fluid and dynamic nature (Fig. 3C). Additionally, treatment with 1,6-hexanediol, a known disruptor of weak hydrophobic interactions in LLPS [46], dissolved the VP4 condensates (Fig. 3D). These results indicate that myristoylated VP4 undergoes LLPS to form biomolecular condensates on the membrane.

**Figure 3:**
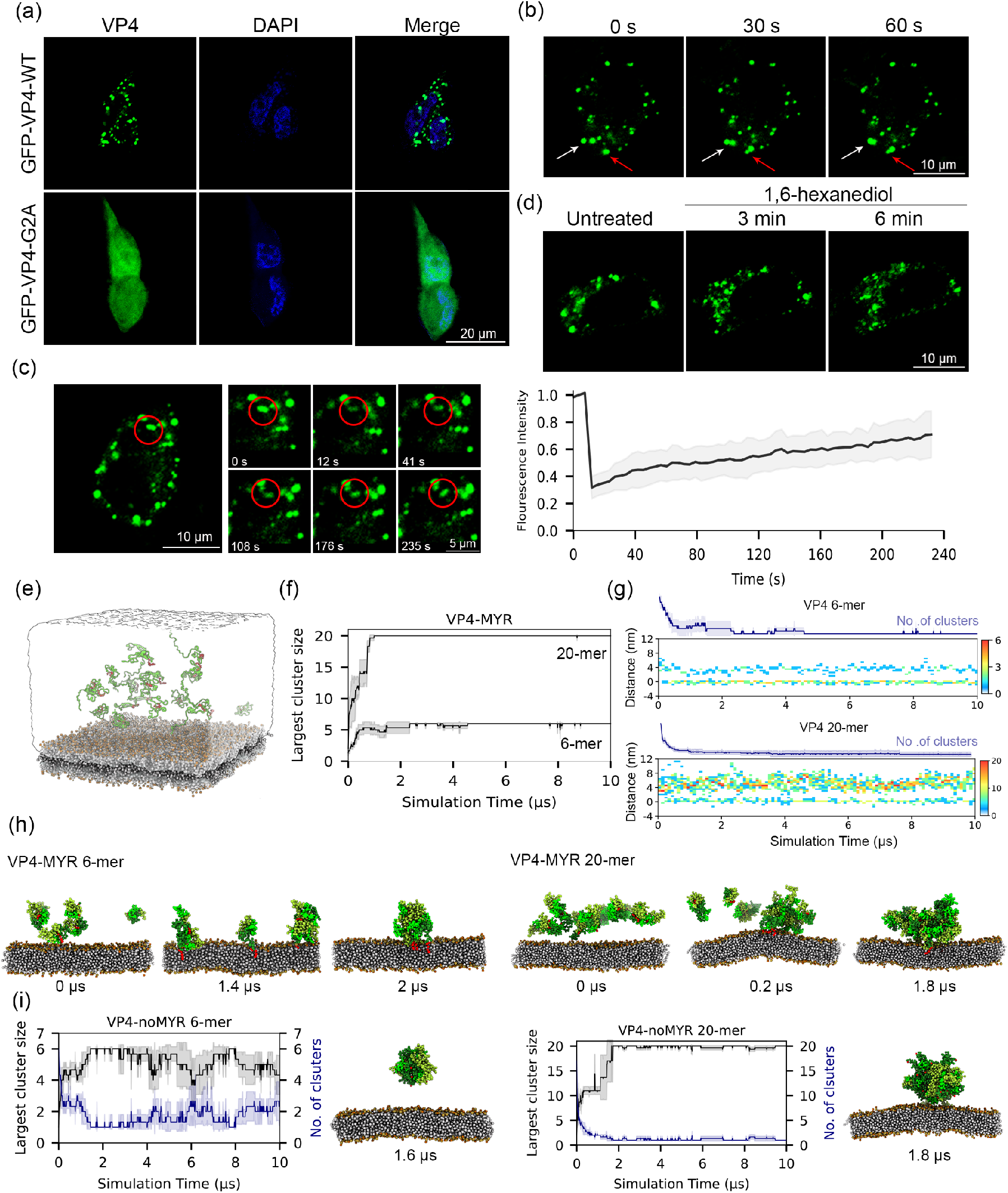
The myristoyl group promotes VP4 condensate formation on membranes. (A) Confocal microscopy images of EGFP-tagged wild-type (WT) VP4 and the non-myristoylated G2A mutant expressed in HEK-293T cells. Scale bar: 20 *µ*m. (B) Time-lapse imaging of VP4-MYR condensate dynamics on a supported lipid bilayer (SLB). Scale bar: 10 *µ*m. (C) FRAP assay quantifying the internal mobility of VP4 condensates on SLBs. Scale bar: 10 *µ*m and 5 *µ*m. (D) Effect of 1,6-hexanediol (10% w/v) treatment on the stability of VP4 condensates. Scale bar: 10 *µ*m. (E) Setup for CGMD simulations investigating VP4 condensation and membrane interaction. Six or twenty myristoylated VP4 proteins were initially placed with a minimum distance of 40 Å from the membrane; three independent 10 *µ*s simulations were performed for each system. (F) Time-dependent analysis of the number of VP4 clusters and the size of the largest cluster during CGMD simulations. (G) Time-dependent distribution of myristoyl moieties along the membrane normal (z-axis). The axis was divided into 0.5 nm bins, and the count of myristoyl groups within each bin is plotted. (H) Representative simulation snapshot illustrating myristoyl-mediated membrane insertion and the nucleation of a VP4 condensate. (I) Control CGMD simulations of non-myristoylated VP4 proteins (three independent 10 *µ*s replicates).

We next used CGMD simulations to explore the molecular basis of this membrane-associated LLPS. Multiple myristoylated VP4 monomers were randomly placed above a membrane bilayer (Fig. 3E). We simulated systems with six and twenty VP4 monomers to investigate the effect of protein density. The six-monomer system was chosen based on reports that CVB3 VP4 can form a six-fold symmetric pore [43], while the twenty-monomer system allowed us to study concentration effects. The time-dependent number of VP4 clusters showed that condensates form in both cases, with aggregation occurring faster and forming more stable condensates in the twenty-monomer system (within 1 *µ*s vs. 2.37 *µ*s) (Fig. 3F). To understand the role of the myristoyl moiety, we calculated its time-dependent distribution along the z-axis, quantifying the synergy between membrane insertion and cluster formation. As shown in Fig. 3G, some myristoyl groups inserted into the membrane shortly after the simulation began, while others remained above the membrane. The membrane-anchored VP4 proteins then appeared to capture the myristoyl groups of other VP4 monomers via their flexible and hydrophobic C-termini, ultimately leading to the formation of stable surface condensates. The myristoyl groups that were not membrane-inserted spontaneously aggregated into a tight hydrophobic core, forming numerous contacts (Fig. 3H-I). In the mature condensate, this myristoyl core was wrapped by other hydrophobic residues from VP4, enhancing inter-residue interactions (Fig. 3H and Fig. S5). This structural analysis suggests a dual role for the myristoyl group: it initiates membrane anchoring and also promotes LLPS. To verify this, we simulated systems with non-myristoylated VP4 proteins (Fig. 3I). The aggregation ability was greatly compromised in both the six- and twenty-monomer cases, showing a trend toward disassembly, in contrast to the stable condensates formed by myristoylated VP4. This is because non-myristoylated VP4 lacks both an anchor to the membrane and a driver to form the hydrophobic core to nucleate the condensate, thereby hindering condensate formation. Similarly, simulations starting from pre-formed VP4 condensates, but with the myristoyl groups removed, led to rapid condensate disassembly (Fig. S6). We also simulated twenty myristoylated VP4 monomers in pure solution (without a membrane) and found that aggregation occurred only after 2 *µ*s, approximately 1 *µ*s later than aggregation in the presence of a membrane. More importantly, fluctuations in the size of the formed aggregates suggest that, even when aggregates formed in solution, they were notably unstable, with dissociation events continuing to occur, in contrast to the consistently stable aggregates formed on the membrane(Fig. S7), indicating that the membrane promotes VP4 condensate formation.

### 3.4 Dynamic condensate facilitates VP4 protein penetration by remodeling the membrane

We next explored the functional implications of the formed VP4 condensate on membrane. We first compared condensates formed by 6 VP4 monomers with those formed by 20 VP4 monomers and found that they exhibit distinct internal protein dynamics. Autocorrelation analysis of the Rg revealed that for the 6-mer condensate, the autocorrelation decayed more slowly at short lag times (*<* 1 *µ*s) than for the 20-mer condensate (Fig. 4A). Since the 6-mer aggregation has a slower relaxation time (Fig. 3F), such autocorrelation difference is unlikely to be due to differences in relaxation rates. Additional autocorrelation analyses starting from the 2.5, 3.0, and 4.0 *µ*s time points of the 10 *µ*s Martini CGMD simulations (Fig. S8) consistently showed that the 20-mer condensate has faster autocorrelation decay at short lag times (*<* 1 *µ*s) than the 6-mer condensate. Thus, the 20-mer condensate undergoes faster large-scale shape fluctuations, indicative of higher internal dynamics.

**Figure 4:**
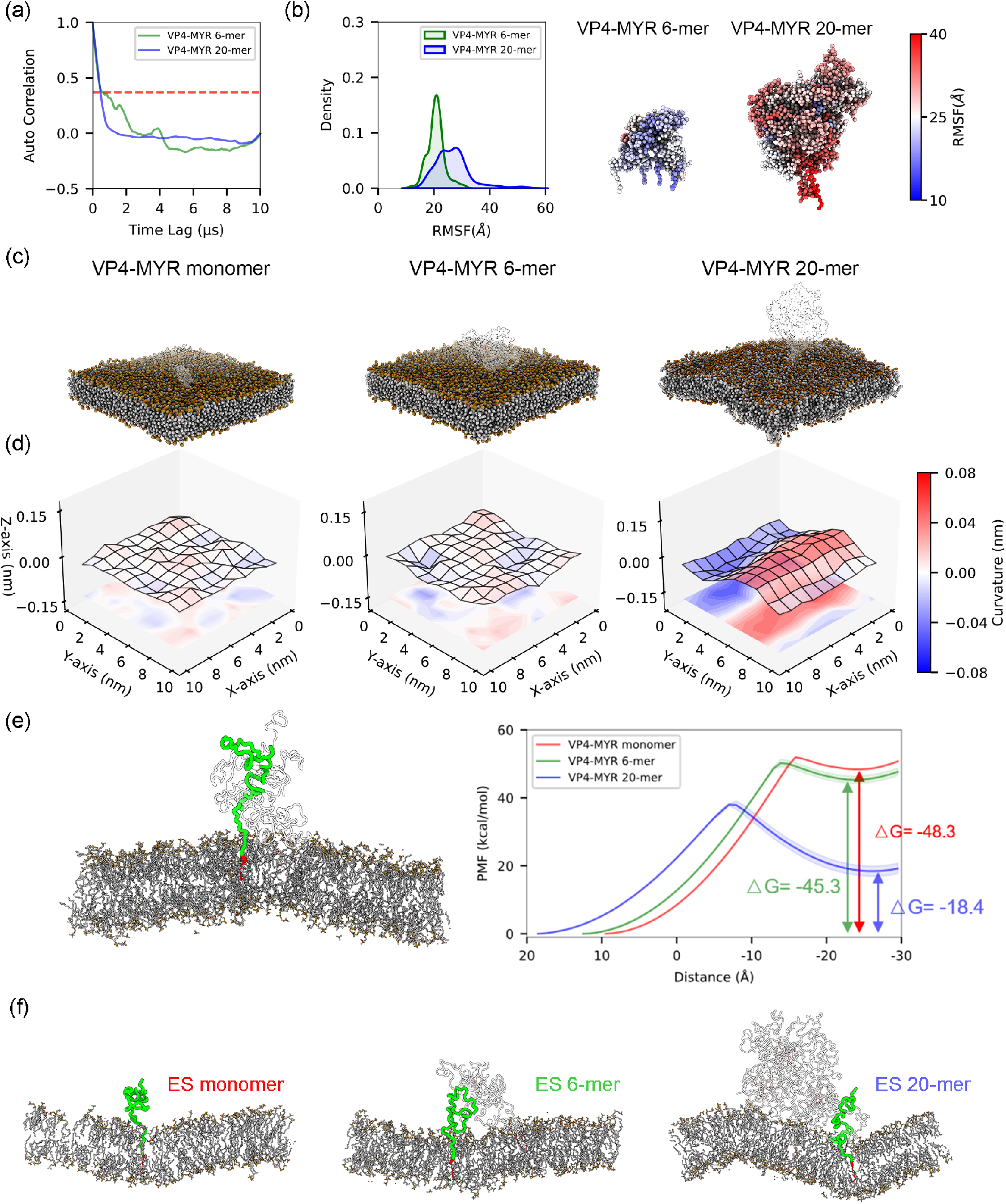
Dynamic VP4 condensates remodel the membrane to facilitate protein penetration. (A) Autocorrelation function of the Rg for the 6-mer and 20-mer VP4 condensates. (B) Distribution of RMSF values for VP4 residues within the 6-mer and 20-mer condensates. A representative condensate structure is shown with residues color-mapped according to their RMSF. (C) Representative simulation snapshots of membrane configurations in the presence of a VP4 monomer, a 6-mer condensate, and a 20-mer condensate. (D) Quantitative analysis of membrane curvature. The local membrane topography is represented by the height function *h*(*x, y*) of the lipid midplane (contour map). Cross-sectional profiles (xz and yz planes) illustrate the membrane deformation induced by the condensates. (E) PMF profiles for the translocation of a VP4 protein from pre-formed condensates into the membrane. The thermodynamically stable end state (ES), characterized by a fully buried myristoyl moiety (red), is labeled. (F) Structural illustrations of the ES configurations for the three systems (monomer, 6-mer, and 20-mer).

We then calculated the backbone RMSF of individual residues within the condensates. The distribution of RMSF values for the 20-mer condensate peaked at 28 Å, which is significantly larger than the peak at 21 Å for the 6-mer (Fig. 4B). Moreover, the RMSF values for the 20-mer spanned a wider range (13 to 56 Å) compared to the 6-mer (11 to 31 Å), demonstrating greater heterogeneity in its internal fluctuations. Mapping residue RMSF values onto a representative condensate snapshot visually confirms this heightened heterogeneity in the 20-mer system. To reveal the origin of this dynamic difference, we performed a time-dependent analysis of residue-residue contacts within the two condensates (Fig. S10). We found that increasing the number of VP4 monomers increases the number of hydrophobic myristoyl cores within the condensate, which compete for residue contacts. This competition weakens the individual interaction strengths within the hydrophobic core network, thereby enhancing the overall fluidity and structural dynamics of the condensate (Fig. S5).

Notably, while the 6-mer condensate exhibited uniformly low RMSF values, the 20-mer condensate showed high dynamics not only in its interior but also for the VP4 molecules directly contacting the membrane (Fig. 4B). We next asked whether this difference in internal dynamics translates to a difference in the ability to remodel the underlying membrane. Structural inspection showed that the membrane beneath the 6-mer VP4 condensate remains overall flat, resembling the membrane with only monomeric VP4 anchored to it (Fig. 4C). In contrast, the 20-mer condensate induced large-scale membrane curvature. Quantitative curvature analysis confirmed that the 20-mer condensate generates membrane regions with high magnitudes of both positive and negative curvature, underscoring its potent membrane-remodeling capability (Fig. 4D)

To directly link membrane remodeling to VP4’s primary function of membrane penetration, we computed the potential of mean force (PMF) for translocating a single VP4 protein from the condensate into the lipid bilayer (Fig. 4E). The translocation of an isolated VP4 monomer without a condensate was the most thermodynamically unfavorable, with an energy barrier of approximately 48.3 kcal/mol (Fig. 4E). Translocation from within the 6-mer condensate slightly reduced this barrier to 45.3 kcal/mol. Remarkably, when starting from the 20-mer condensate, the translocation energy barrier was dramatically reduced to about 18.4 kcal/mol. The heights of these energy barriers are consistent with the membrane curvature profiles. As a critical control, we recalculated the PMF for the 20-mer system while applying a external pressure to maintain a flat membrane; under these conditions, the energy barrier increased from 18.4 kcal/mol to 80.9 kcal/mol (Fig. S11). These data demonstrate that the 20-mer VP4 condensate facilitates the essential translocation process by remodeling the membrane, thereby significantly lowering the energy barrier. Furthermore, in the 20-mer case, the transmembrane end-state (ES) is substantially more stable, by 27-30 kcal/mol, than the end-states in the other two systems (Fig. 4E). Structural inspection of the ES (Fig. 4F) provides a plausible explanation for this stability: a curved membrane is a common feature allowing VP4 burial in all ES structures. The pre-remodeled membrane beneath the 20-mer condensate naturally provides this optimal curved environment, pre-configuring the membrane for favorable VP4 insertion.

### 3.5 Potential role of myristoylation in stabilizing a pore formed by VP4 proteins

Our data have established that myristoylation-driven condensation of VP4 on the membrane facilitates protein penetration by lowering the associated energy barrier. While the ability of picornaviral VP4 proteins to form size-selective membrane pores is well-documented [5, 43, 47], the atomic-level structure of these pores remains unknown. Given that the pore-forming capability has been mapped to the N-terminal region of VP4 [43], we sought to explore a plausible molecular mechanism: could the functional end-state of penetration involve the formation of a structured, multimeric pore, and if so, how might myristoylation contribute ?

As *α*-helices are common structural motifs in membrane-active peptides [48] and have been implicated in the membrane interaction of other viral proteins [47], we first investigated whether disordered VP4 could spontaneously folds into helix in membrane. We focused on the N-terminal 20 residues and performed metadynamics simulations to explore the folding profile. In water, both the unmodified and myristoylated VP4 N-termini remained largely disordered (stable *α*-helicity is around 0.2, Fig. 5A). However, within a POPC bilayer, a folded state with *α*-helical content *>* 0.6 is favored (Fig. 5A). Notably, the presence of the myristoyl group further promoted folding, with the modified peptide (VP4-MYR) sampling states with a higher maximum helicity (∼0.8) compared to the unmodified counterpart (∼0.68). This suggests that the hydrophobic environment of the membrane, particularly when combined with the lipid anchor, induces and stabilizes helical structure in the VP4 N-terminus.

**Figure 5:**
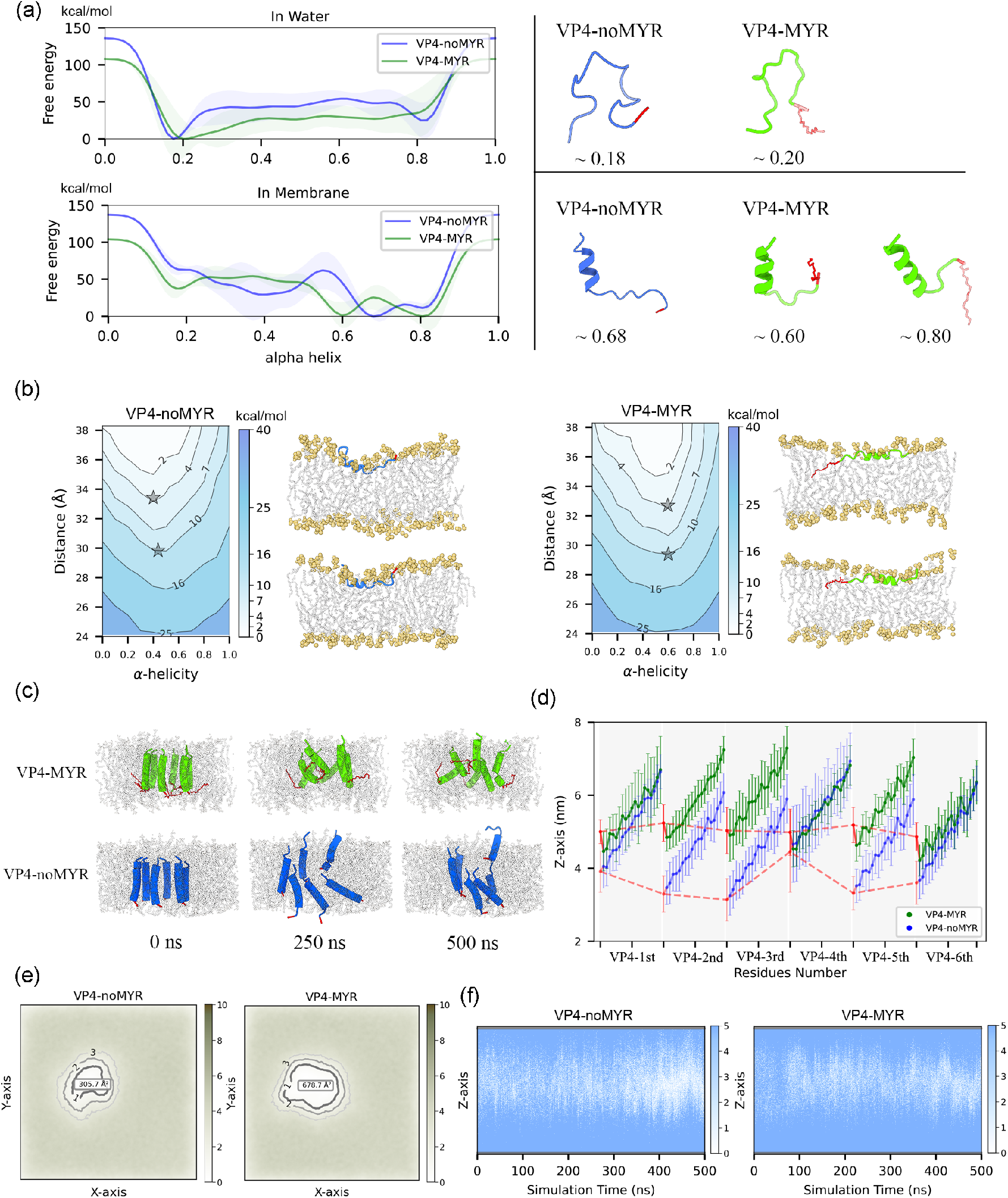
Myristoylation stabilizes a multimeric pore formed by VP4 N-terminal helices. (a) Free energy profile of the coil-to-helix transition for the N-terminal 20 residues of VP4 in aqueous and lipid environments, with and without myristoylation (MYR), respectively. (b) Two-dimensional free energy landscape from umbrella sampling, showing the relationship between the helicity propensity and the membrane embedding depth of the VP4 N-terminus. Asterisks mark representative conformations at the 4 and 10 kcal/mol free energy contours. (c) Structural evolution of a model pore formed by six VP4 N-terminal helices. The initial configuration featured six vertically arranged helices, which were subsequently refined via three independent 500 ns MD simulations. Snapshots from one representative simulation are shown. (d) Range of residue Z-coordinates sampled during the MD simulations. Error bars represent the standard deviation across the three replicates. (e) Lipid density projected onto the XY plane from one 500 ns simulation. Data from the remaining two runs are shown in Fig. S12. (f) Time-dependent number of water molecules within a cylindrical volume encompassing the pore, projected along the Z-axis (n=3 independent runs). Data from the remaining two runs are shown in Fig. S13.

We next explored whether this helical-switching propensity correlates with the embedding depth of the N-terminus within membrane, using two-dimensional PMF calculations. As shown in Fig. 5B, we found that for both unmodified and myristoylated VP4 N-terminus, folding into an *α*-helix is thermodynamically coupled with membrane insertion. However, the myristoyl modification confers a distinct thermodynamic advantage, allowing the VP4 N-terminus to embed deeper into the membrane while maintaining a higher degree of helicity. This is illustrated by comparing two representative snapshots at the 4 and 10 kcal/mol free energy contours (starred points on the potential surface). Both Myr-modified and unmodified VP4 fragments in these representative conformations have been completely inserted into the lipid bilayer. However, at the same free energy cost of 10 kcal/mol, the myristoylated VP4 could penetrate approximately 1.2 Å deeper into the membrane while maintaining a helicity (*α* ∼0.6) significantly greater than that of the unmodified peptide (*α* ∼0.4) (Fig. 5B). This indicates that myristoylation not only promotes helix formation but also helps stabilize the helical conformation within the membrane interior. It is important to note that a complete translocation remained energetically prohibitive for both peptides, consistent with a pore-forming, rather than a translocating, mechanism.

Finally, we conducted all-atom MD simulations to investigate whether a pore formed by VP4 helices would be plausible within the membrane. Guided by TEM imagery suggesting six-fold symmetry [5], we inserted six helical VP4 N-termini into the lipid bilayer to construct a pore model and performed three independent 0.5 *µ*s simulations to assess its stability. As shown in Fig. 5C, in the myristoylated (MYR) system, the myristoyl moieties anchored the helices via hydrophobic interactions with lipids, maintaining a relatively stable pore architecture. In contrast, the non-myristoylated counterpart exhibited significant fluctuation and a tendency to disassemble (Fig. 5C,D). Consequently, the unmodified pore was susceptible to lipid occlusion, resulting in a smaller solvent-accessible cavity and reduced water transport compared to the myristoylated pore (Fig. 5EF).

Thus, our computational data suggests that myristoylation promotes *α*-helical folding, facilitates deep membrane insertion, and stabilizes the supramolecular assembly of the functional pore. This model provides a testable hypothesis linking initial viral condensation to the molecular mechanism of membrane disruption.

## 4 Conclusions

Membrane breaching is a vital step for viruses to deliver their genomic material into host cells to establish infection. Non-enveloped viruses rely on the externalized small VP4 or VP4-like proteins from their capsid to interact with and breach the membrane. This multi-step process has been understood only at a phenotypic level, with no clear structural mechanism revealed. This mystery is particularly intriguing because some VP4 proteins are intrinsically disordered, and this disordered nature requires myristoylation to license their membrane-breaching function. In this study, by combining multiscale computational modeling with experimental assays, we reveal the structural mechanism underlying the myristoylation-dependence of the disordered VP4 protein from CVB3 (Fig. 6). We demonstrate that myristoylations canonical function of conferring membrane-targeting ability represents a mutual selection for VP4: myristoylation allows VP4 to anchor to the membrane, yet this ability requires VP4 to be disordered. Furthermore, we show that myristoylation drives the disordered VP4 proteins to undergo phase separation on the membrane surface, a process that remodels the underlying membrane to lower the energy barrier for VP4 insertion. Lastly, we discovered a disorder-to-helix transition in the N-terminus of VP4 upon membrane embedding and demonstrated that myristoylation enhances this helical folding propensity. It also plays a stabilizing role in a putative transmembrane pore formed by these VP4 helices. Our findings uncover a mechanism wherein myristoylation licenses a disordered viral protein for membrane penetration, acting not merely as a simple anchor but as a key driver of functional, collective behavior through phase separation. This work provides a unifying structural and biophysical framework to explain the long-standing mystery of myristoylation-dependence in viral membrane-penetrating proteins.

**Figure 6:**
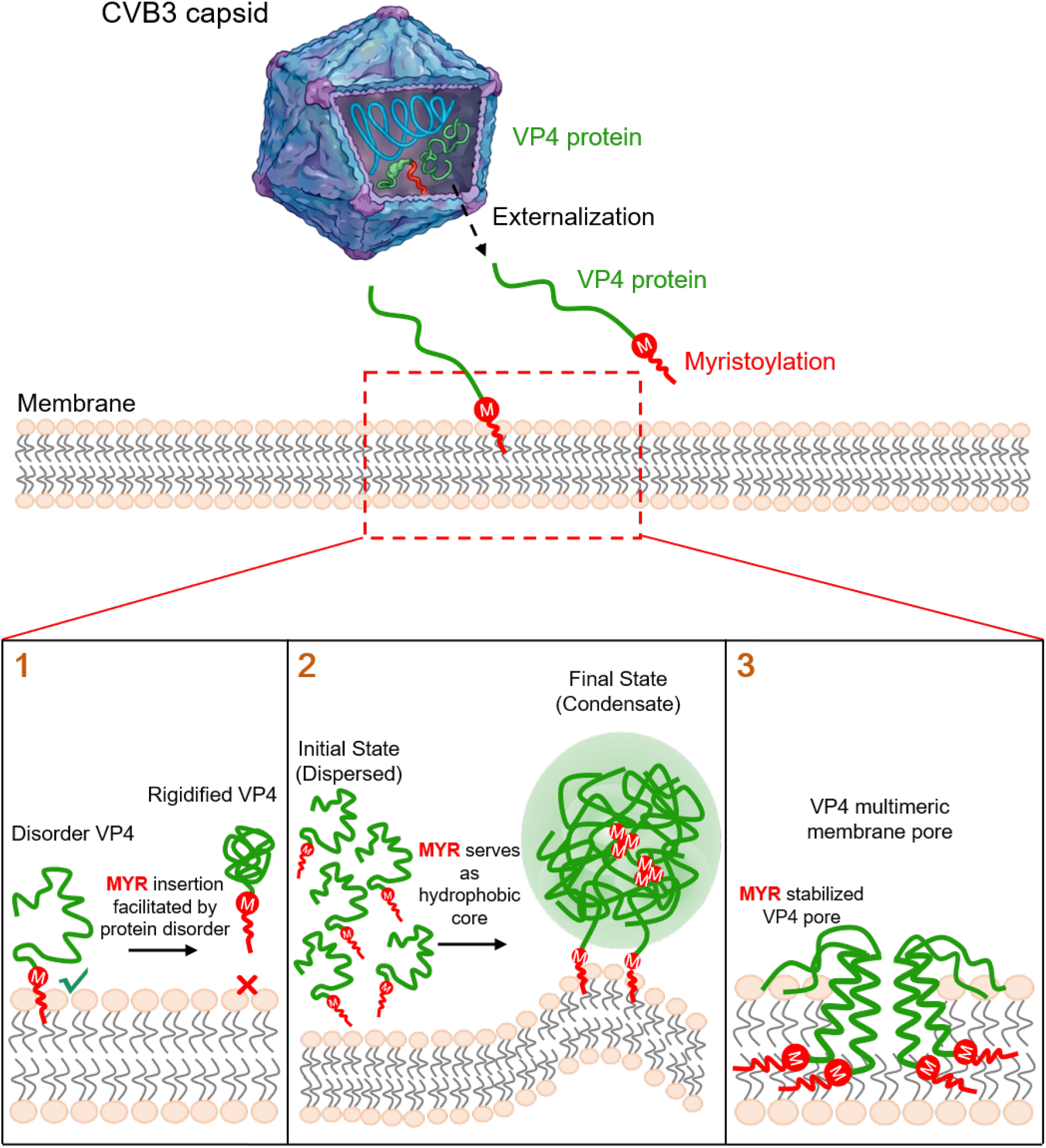
The structural basis for myristoylation’s multi-faceted role in enabling the disordered VP4 protein to breach the host cell membrane.

## 5 Acknowledgements

Research reported in this work was supported by the National Natural Science Foundation of China (32501103), Natural Science Foundation of Heilongjiang Province (PL2024B022), Heilongjiang Provincial Postdoctoral Science Foundation (LBH-Z24211), and China Postdoctoral Science Foundation (2023M730827, 2025MD784111).

## 5.1 Data Availability Statements

Molecular dynamics simulations were performed using the Amber20 and GROMACS 2020/22 package, with detailed protocol reported in the method section of the manuscript. Plotting was performed using Matplotlib, and structural rendering was done with ChimeraX. Scripts used for this project are available at: https://github.com/bsu233/bslab/tree/main/2025VP4. Simulation trajectories are available upon request.

Supplementary Information (SI):

---

**Figure S1:**
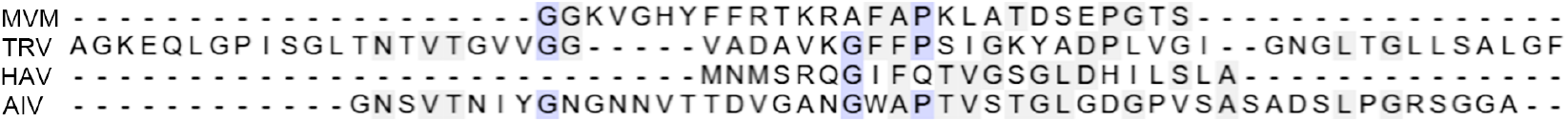
Sequence alignment of VP4 proteins that do not need myristoylation modification to function.

**Figure S2:**
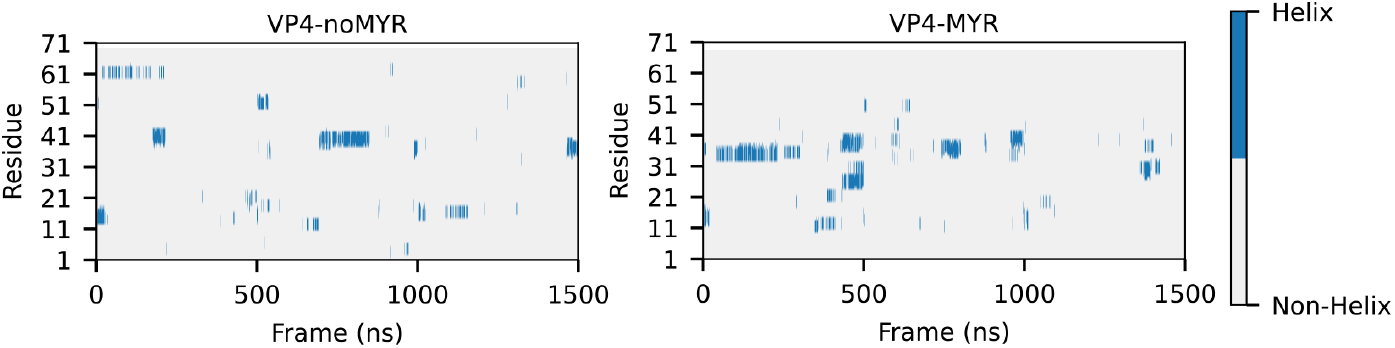
Per-residue helicity measured by DSSP from atomistic 1500 ns MD simulations of myristoylated and non-myristoylated CVB VP4 proteins.

**Figure S3:**
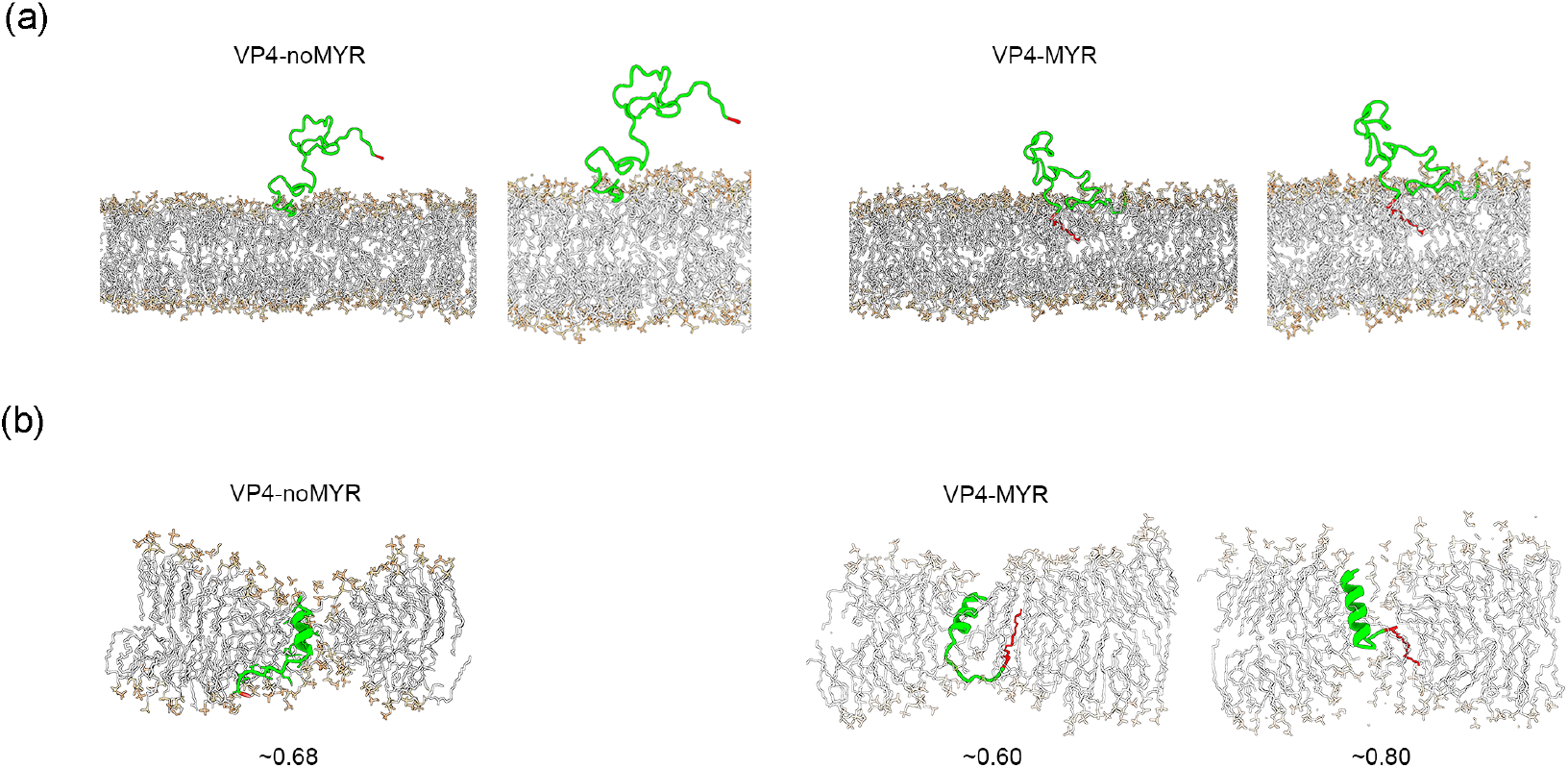
Atomistic VP4 structures. (a) Backmapped atomistic structure of a Martini CGMD-sampled, membrane-bound VP4 peptide. (b) Favorable helical conformations of a VP4 fragment (N-terminal 20 residues) fully embedded in a POPC lipid bilayer, as sampled by WT-MetaD simulations. Helicity values are labeled.

**Figure S4:**
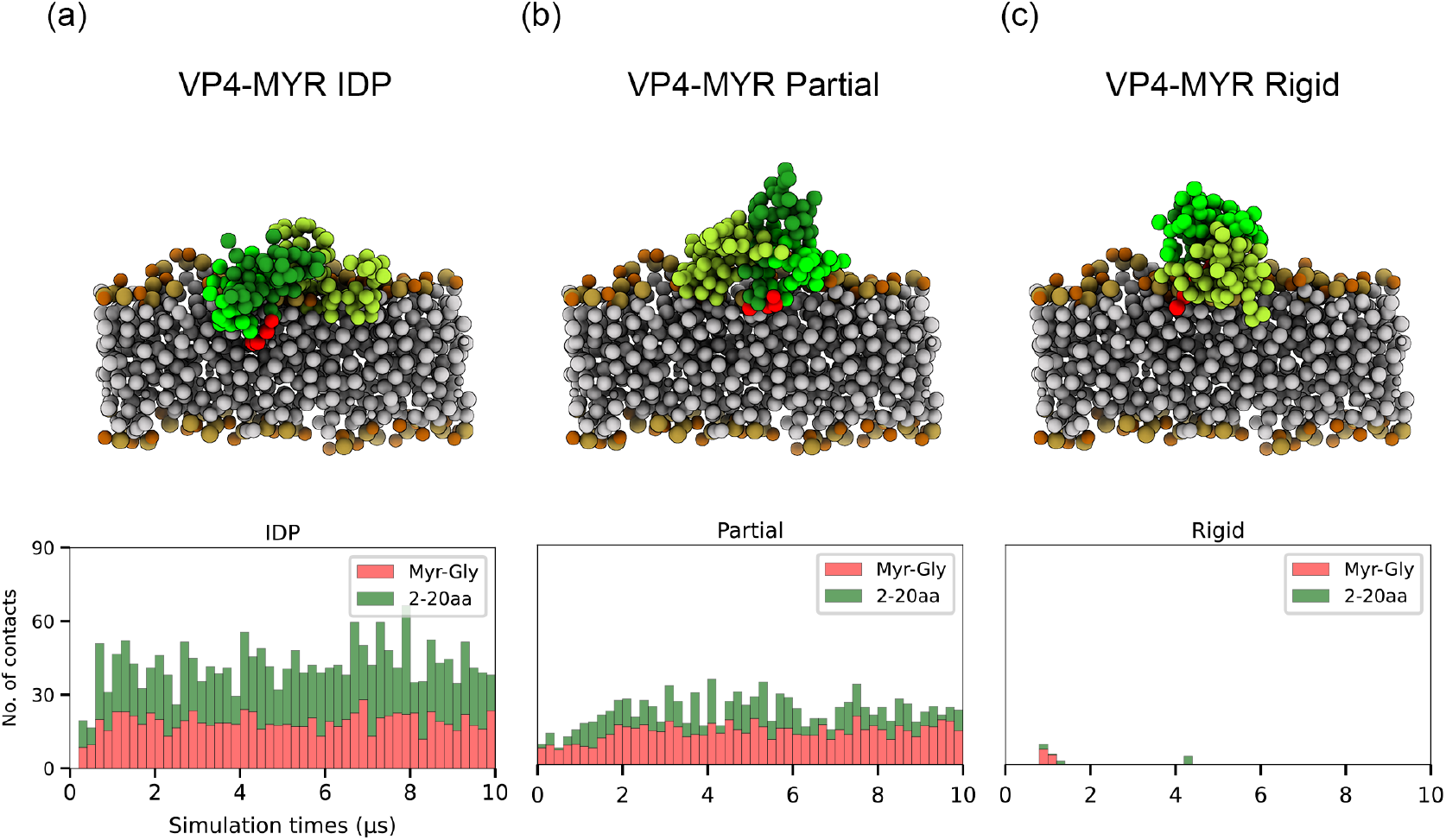
MD simulations of reduced flexibility VP4 binding to membrane. Panel (a) shows the Martini coarse-grained molecular dynamics (CGMD) sampled bound states of native, disordered VP4 with the membrane. Panels (b) and (c) show the corresponding states for a VP4 protein with partially reduced flexibility and a rigid VP4 protein, respectively. The N-terminal myristoylation moiety is colored red. The bar plot at the bottom shows the time-dependent number of contacts between the membrane and the myristoylated GLY residue (orange) and between the membrane and the first 20 residues of VP4 (green).

**Figure S5:**
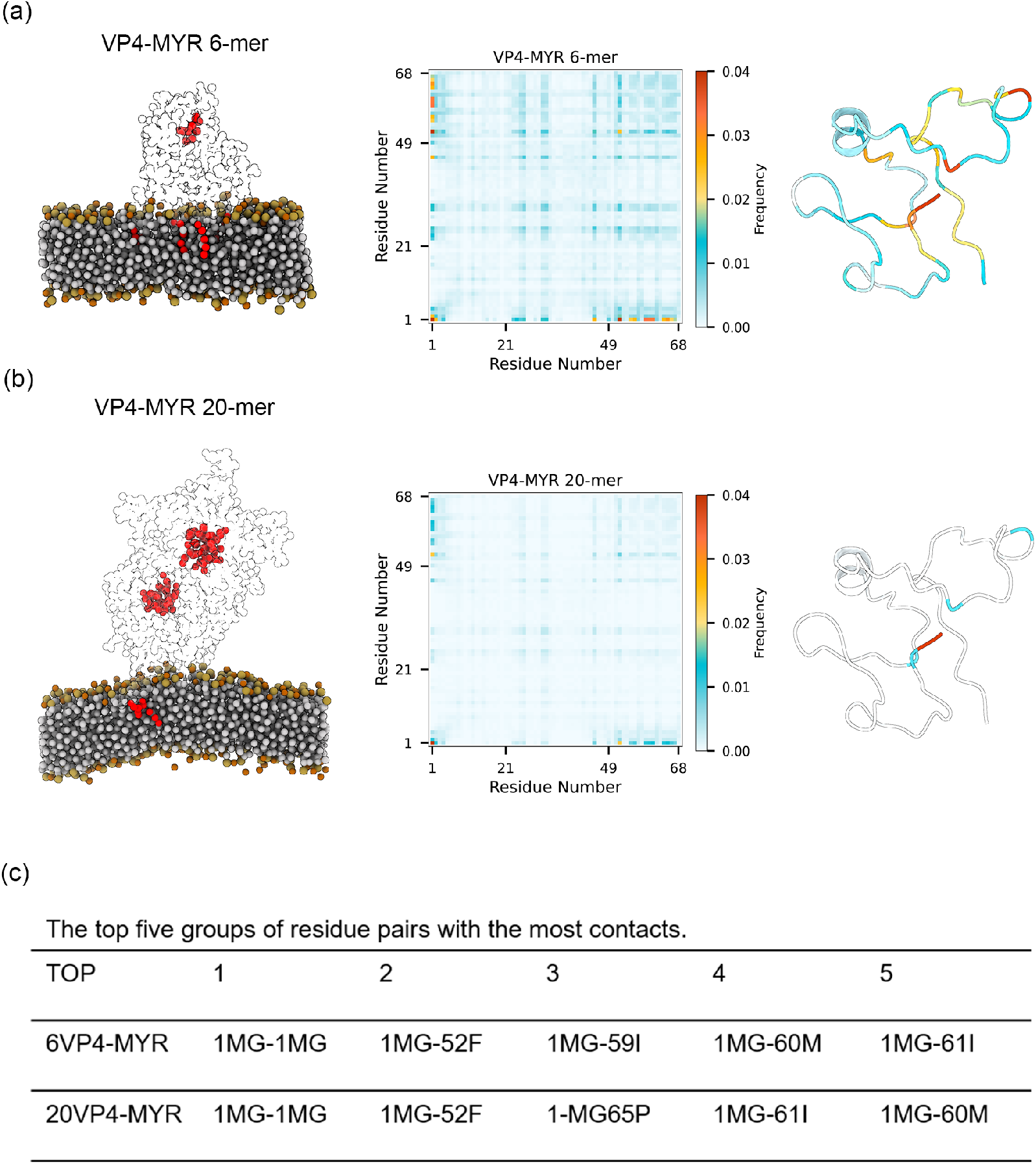
The myristoylated GLY residues form hydrophobic cores within the condensate. (a) A snapshot of the final 6-mer condensate formed on the membrane, with the myristoylation moiety colored red. The heatmap shows the average residue-residue contact number within the 6-mer condensate. The right-most panel shows the average contact number mapped onto a single VP4 protein structure. (b) The same analysis as in panel (a) for the 20-mer condensate. (c) The top five residue pairs with the highest contact frequencies. “MG” refers to the myristoylated glycine residue.

**Figure S6:**
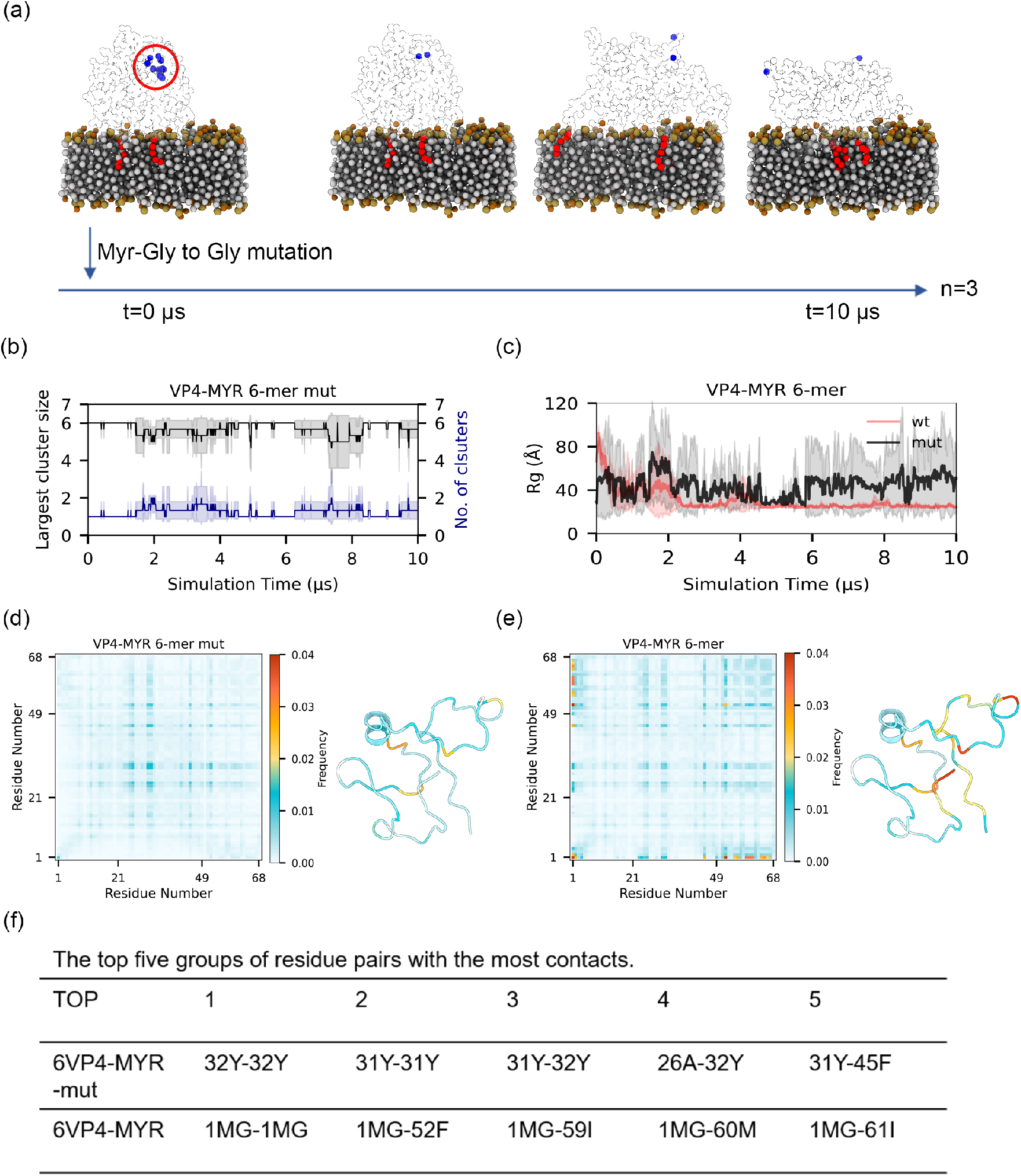
Effects of annulling myristoylation within the formed VP4 condensate. (a) Schematic of the simulation strategy: myristoylation sites in a pre-formed 6-mer VP4 condensate were mutated to glycine, followed by molecular dynamics (MD) investigation. (b) Time evolution of the number of VP4 clusters and the size of the largest cluster after the mutation. (c) Radius of gyration (Rg) of the condensate following the mutations. (d, e) Average residue-residue contact numbers between VP4 residues in the wild-type condensate (d) and the mutant condensate (e). These contact numbers are also mapped onto single VP4 protein structures. (f) Top five residue pairs with the highest contact numbers in the wild-type condensate and the mutant condensate, respectively.

**Figure S7:**
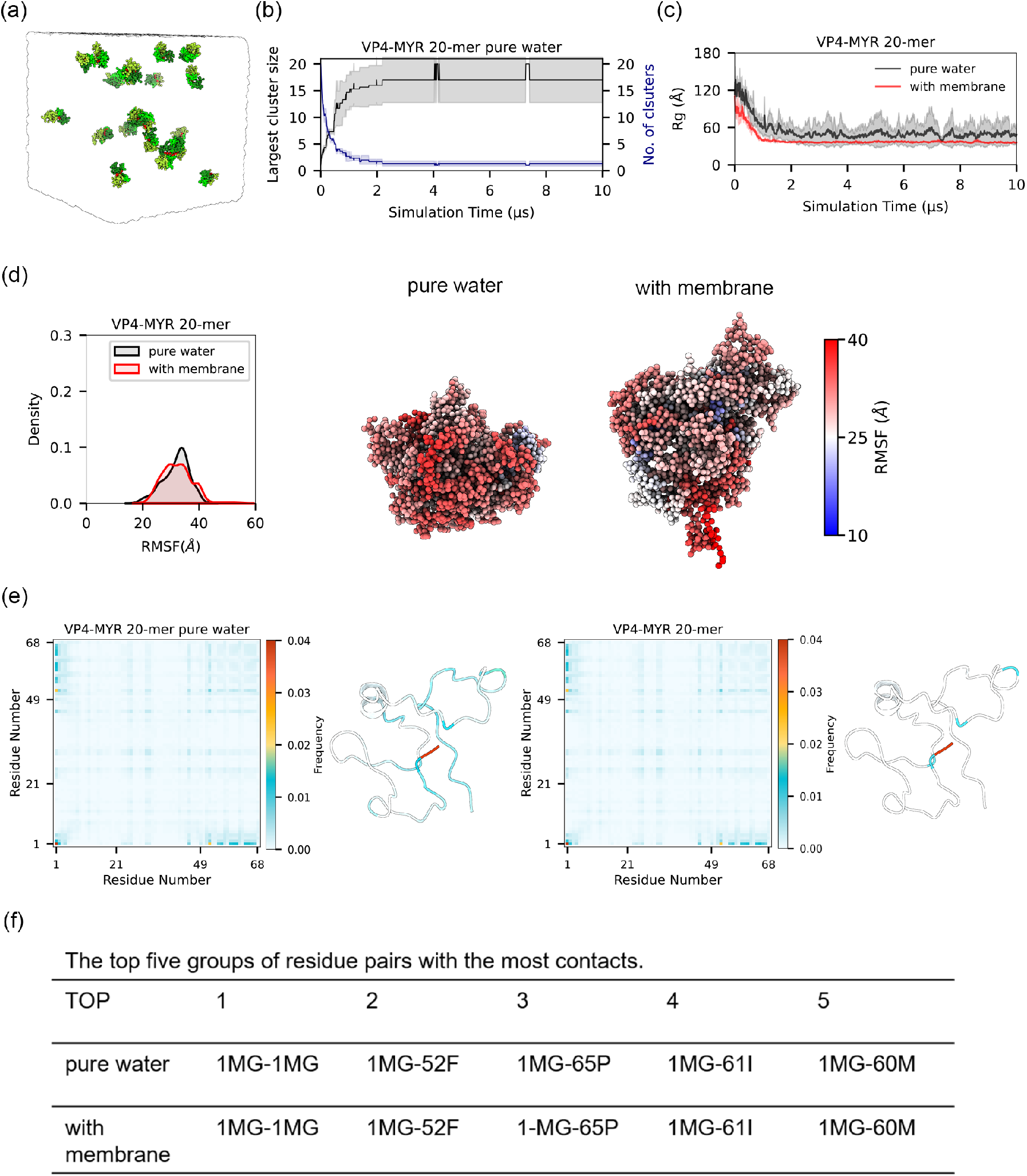
Condensation behavior of VP4-MYR monomer in pure aqueous solution. (a) Schematic of Martini CGMD simulations with 20 VP4-MYR monomers in pure aqueous solution (3 independent runs of 10 *µ*s each). (b) Time evolution of the number of VP4 clusters and the size of the largest cluster in solution. (c) Radius of gyration (Rg) of condensates formed in solution compared to those formed on the membrane. (d) Distribution of RMSF values calculated from VP4 condensates formed in solution versus those formed on the membrane. A representative condensate structure is shown, with residues color-mapped according to their RMSF values. (e) Average residue-residue contact numbers calculated from VP4 condensates formed in solution and on the membrane. (f) Top five residue pairs with the highest contact frequencies in condensates formed in solution versus those formed on the membrane.

**Figure S8:**
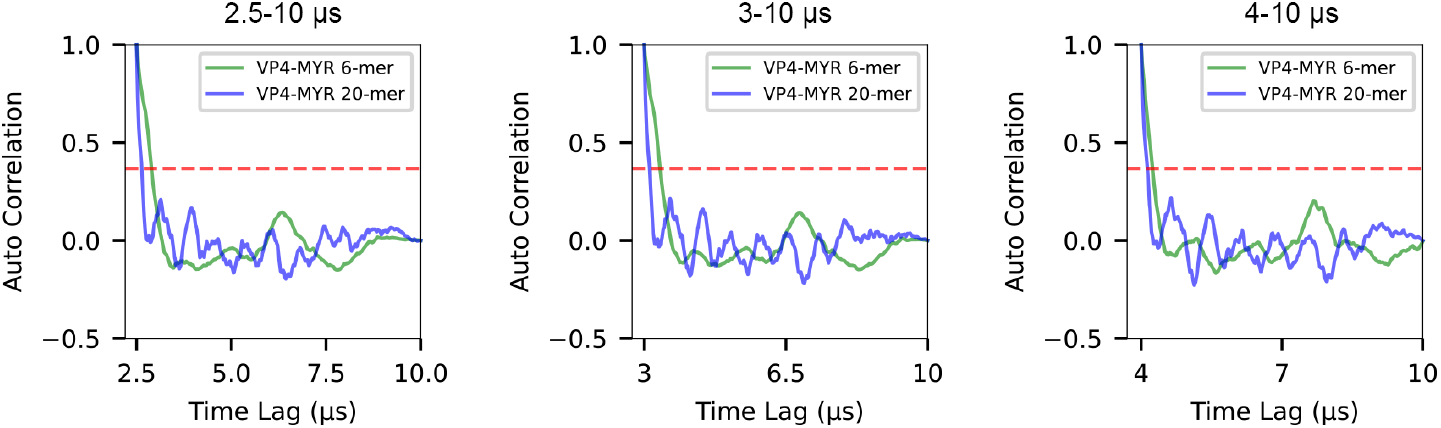
Autocorrelation analyses of the condensate radius of gyration (*R_g_*) initiated from the 2.5, 3.0, and 4.0 *µ*s time points of the 10 *µ*s Martini CGMD simulations.

**Figure S9:**
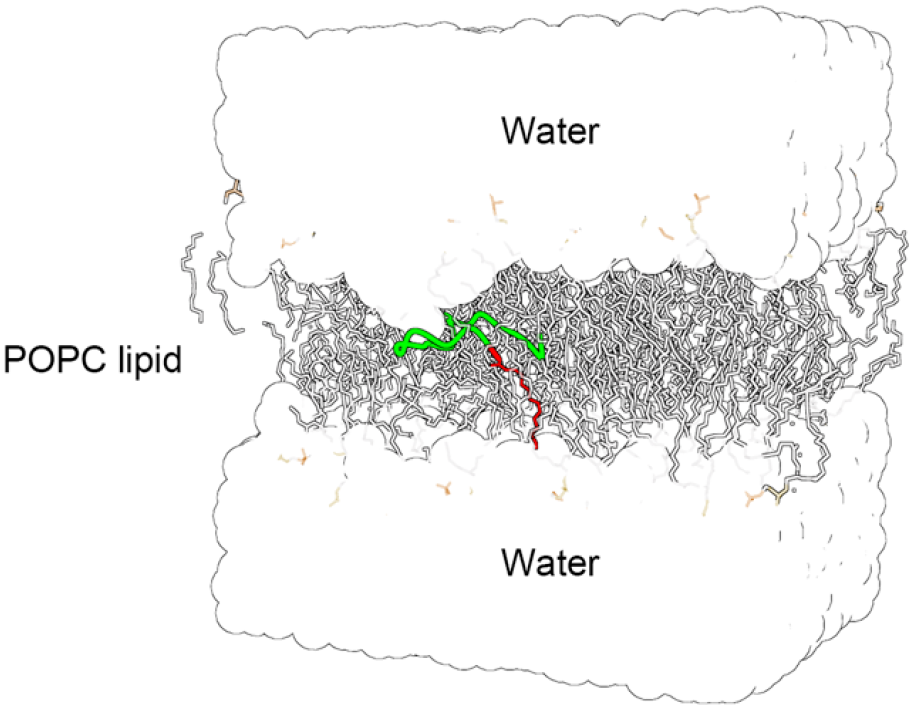
Starting structure for the WT-MetaD simulation exploring the free-energy profile of the VP4 fragment (N-terminal 20 residues) with MYR modification. The VP4 fragment was placed manually at the center of the lipid bilayer. The corresponding no-MYR WT-MetaD simulation used the same starting structure with the MYR moiety removed.

**Figure S10:**
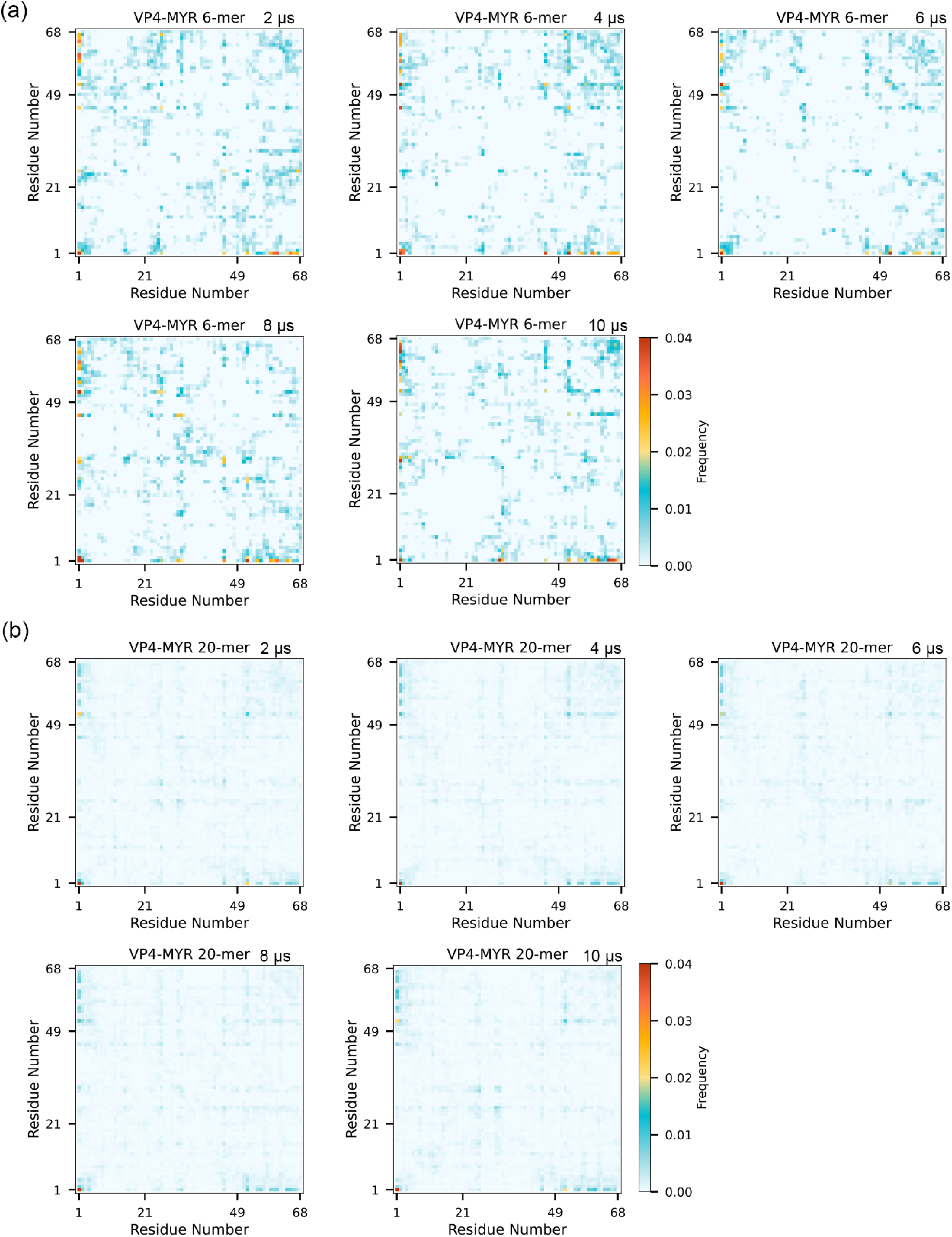
Time-dependent analysis of residue-residue contacts within the condensate. Average residue-residue contact numbers calculated from the VP4-MYR condensates formed on the membrane at different time points (2, 4, 6, 8, and 10 *µ*s). (a) Results for the 6-mer condensate. (b) Results for the 20-mer condensate.

**Figure S11:**
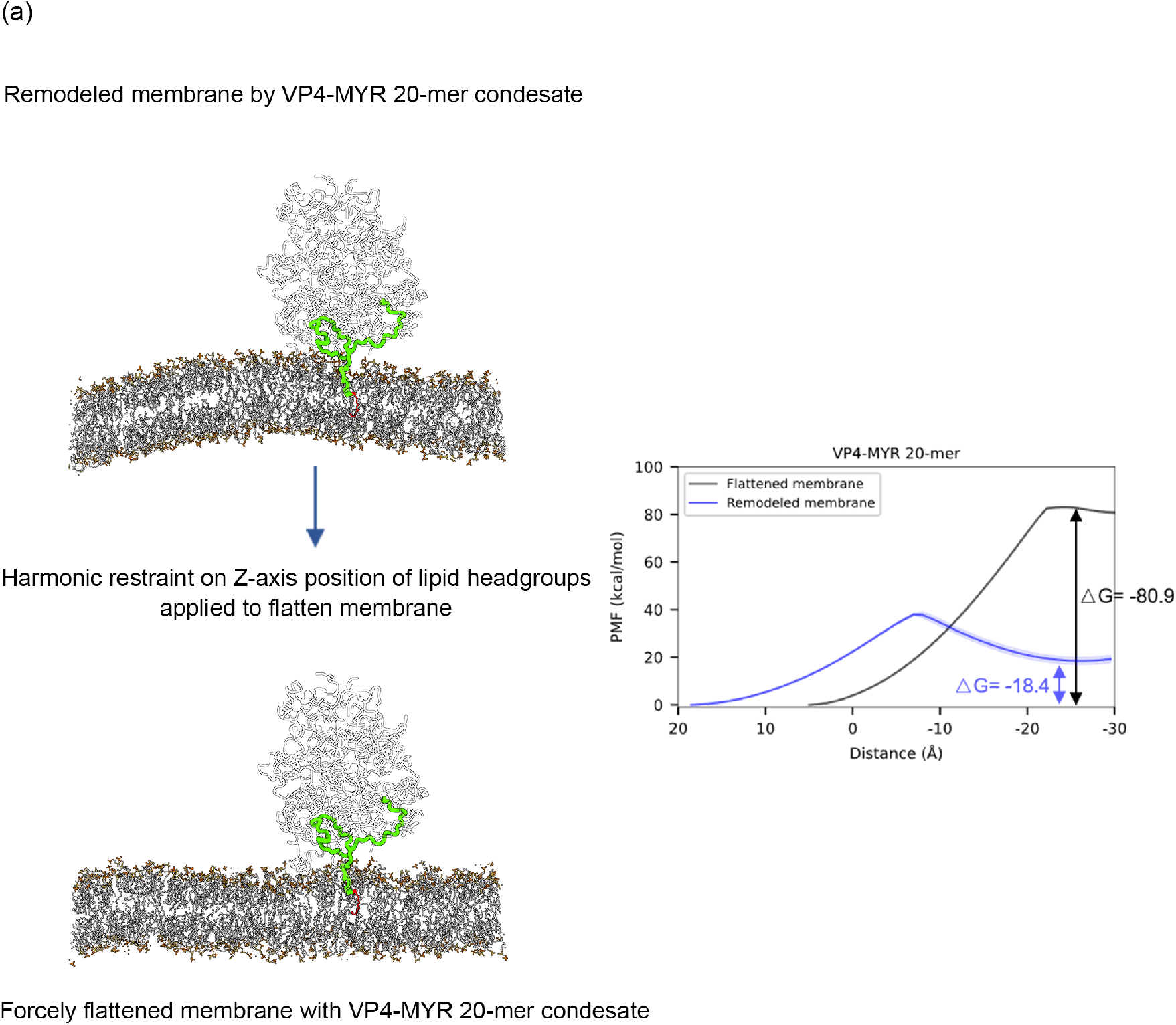
Energetics of VP4 penetrating an artificially flattened membrane. (a) Schematic illustrating the harmonic restraint along Z-axis applied to lipid headgroups to flatten a membrane that had been remodeled by a 20-mer VP4-MYR condensate. (b) Potential of mean force (PMF) profile along the translocation reaction coordinate for a single VP4 molecule penetrating the artificially flattened membrane, compared to the unflattened membrane.

**Figure S12:**
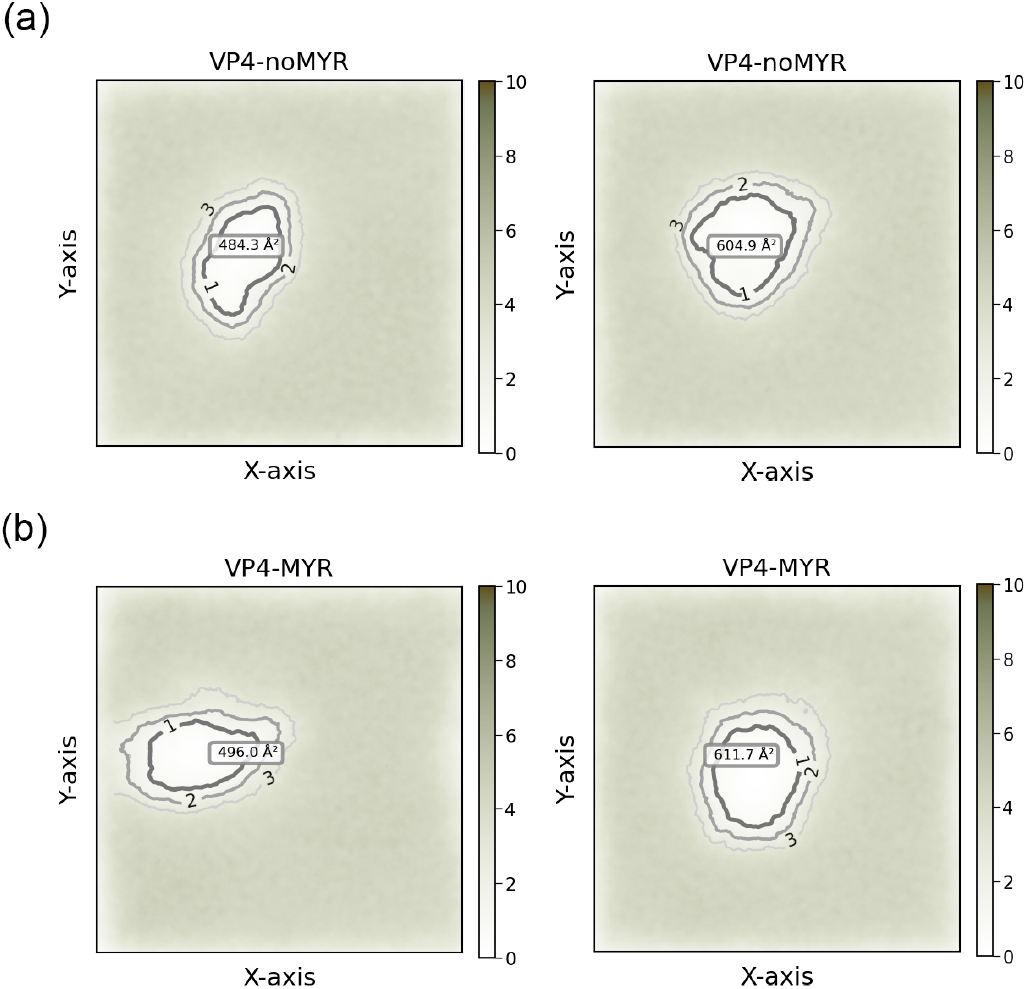
Lipid density profiles across the membrane were calculated from two 0.5 *µ*s all-atom molecular dynamics (MD) simulations investigating the stability of the pore formed by six N-terminal VP4 helices. Results are shown for the system (a) without and (b) with the MYR (myristoylation) modification.

**Figure S13:**
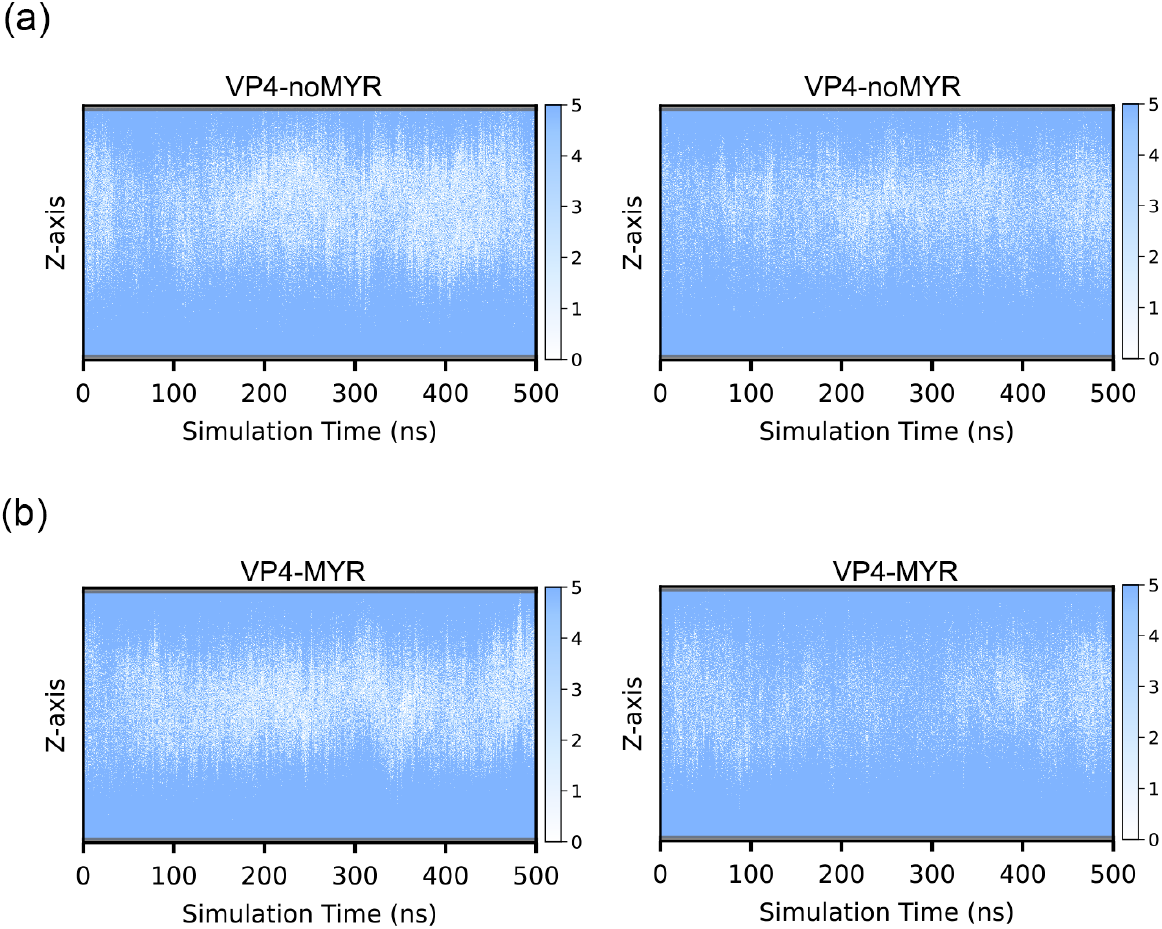
The time-dependent count of water molecules within the pores was analyzed from two 0.5 *µ*s all-atom MD simulations. The distribution of water molecules along the membrane normal (Z-axis) is plotted for the system (a) without and (b) with the MYR (myristoylation) modification.

**Table S1:** Molecular dynamics simulations performed in this study.

| Simulation Types | Cases | AA/CG model | Simulation Length |
| --- | --- | --- | --- |
| **Conventional MD** | VP4-noMYR in solution | AA (Amber20) | 1 μs x 3 |
| | VP4-MYR in solution | AA | 1 μs x 3 |
| | VP4-noMYR with membrane | CG (Gromacs 2020.4) | 10 μs x 3 |
| | VP4-MYR with membrane | CG | 10 μs x 3 |
| | VP4-MYR Partial model with membrane | CG | 10 μs x 3 |
| | VP4-MYR Rigid model with membrane | CG | 10 μs x 3 |
| | VP4-MYR 6-mer with membrane | CG | 10 μs x 3 |
| | VP4-MYR 20-mer with membrane | CG | 10 μs x 3 |
| | VP4-MYR 6-mer mutation with membrane | CG | 10 μs x 3 |
| | VP4-MYR 20-mer in solution | CG | 10 μs x 3 |
| | VP4-MYR 6-mer with membrane | CG | 10 μs x 3 |
| | VP4-noMYR 6-mer N-ter in membrane | AA | 0.5 μs x 3 |
| | VP4-MYR 6-mer N-ter in membrane | AA | 0.5 μs x 3 |
| **Potential Mean Force** | VP4-MYR with membrane | CG (Gromacs 2022.5) | 5.5 μs x 3 |
| | VP4-MYR 6-mer with membrane | CG | 6.4 μs x 3 |
| | VP4-MYR 20-mer with membrane | CG | 5.1 μs x 3 |
| | VP4-MYR 20-mer with flattened membrane | CG | 3.6 μs x 3 |
| **2D Potential Mean Force** | VP4-noMYR with membrane | AA (Gromacs 2022.5) | 3.63 μs |
| | VP4-MYR with membrane | AA | 3.63 μs |
| **MetaDynamics** | VP4-noMYR N-ter in solution | AA (Gromacs 2022.5) | 0.5 μs x 3 |
| | VP4-MYR N-ter in solution | AA | 0.5 μs x 3 |
| | VP4-noMYR N-ter with membrane | AA | 0.5 μs x 3 |
| | VP4-MYR N-ter with membrane | AA | 0.5 μs x 3 |

## References

[1] Margaret Kielian and Félix A Rey. “Virus membrane-fusion proteins: more than one way to make a hairpin”. Nature Reviews Microbiology 4 (2006), 67–76.

[2] Donia Bouzid, Oshra Hadad, Mélanie Bertine, Nadhira Houhou-Fidouh, Audrey Mirand, Xavier Duval, Vincent Bunel, Raphael Borie, Jean Christophe Lucet, Diane Descamps, et al. “Rhinoviruses: Molecular diversity and clinical characteristics”. International Journal of Infectious Diseases 118 (2022), 144–149.

[3] Qiong Liu, Yinpan Shang, Ziwei Tao, Xuan Li, Lu Shen, Hanchi Zhang, Zhili Liu, Zhirong Rao, Xiaomin Yu, Yanli Cao, et al. “Coxsackievirus group B3 regulates ASS1-mediated metabolic reprogramming and promotes macrophage inflammatory polarization in viral myocarditis”. Journal of Virology 98 (2024), e00805–24.

[4] Glyn Stanway. “Structure, function and evolution of picornaviruses”. Journal of General Virology 71 (1990), 2483–2501.

[5] Anusha Panjwani, Mike Strauss, Sarah Gold, Hannah Wenham, Terry Jackson, James J Chou, David J Rowlands, Nicola J Stonehouse, James M Hogle, and Tobias J Tuthill. “Capsid protein VP4 of human rhinovirus induces membrane permeability by the formation of a size-selective multimeric pore”. PLoS pathogens 10 (2014), e1004294.

[6] Pranav NM Shah, David J Filman, Krishanthi S Karunatilaka, Emma L Hesketh, Elisabetta Groppelli, Mike Strauss, and James M Hogle. “Cryo-EM structures reveal two distinct conformational states in a picornavirus cell entry intermediate”. PLoS pathogens 16 (2020), e1008920.

[7] Stephen Curry, Marie Chow, and James M Hogle. “The poliovirus 135S particle is infectious”. Journal of virology 70 (1996), 7125–7131.

[8] Yan Huang, James M Hogle, and Marie Chow. “Is the 135S poliovirus particle an intermediate during cell entry?” Journal of Virology 74 (2000), 8757–8761.

[9] Meng Yuan, Zi-han Song, Mei-dan Ying, Hong Zhu, Qiao-jun He, Bo Yang, and Ji Cao. “N-myristoylation: from cell biology to translational medicine”. Acta Pharmacologica Sinica 41 (2020), 1005–1015.

[10] Jiaming Cao, Meng Qu, Hongtao Liu, Xuan Wan, Fang Li, Ali Hou, Yan Zhou, Bo Sun, Linjun Cai, Weiheng Su, et al. “Myristoylation of EV71 VP4 is essential for infectivity and interaction with membrane structure”. Virologica Sinica 35 (2020), 599–613.

[11] Irena Corbic Ramljak, Julia Stanger, Antonio Real-Hohn, Dominik Dreier, Laurin Wimmer, Monika Redlberger-Fritz, Wolfgang Fischl, Karin Klingel, Marko D Mihovilovic, Dieter Blaas, et al. “Cellular N-myristoyltransferases play a crucial picornavirus genus-specific role in viral assembly, virion maturation, and infectivity”. PLoS Pathogens 14 (2018), e1007203.

[12] Ashutosh Shukla, Aditya K Padhi, James Gomes, and Manidipa Banerjee. “The VP4 peptide of hepatitis A virus ruptures membranes through formation of discrete pores”. Journal of virology 88 (2014), 12409–12421.

[13] James T Kelly, Jessica Swanson, Joseph Newman, Elisabetta Groppelli, Nicola J Stonehouse, and Tobias J Tuthill. “Membrane interactions and uncoating of aichi virus, a picornavirus that lacks a VP4”. Journal of Virology 96 (2022), e00082–22.

[14] Shuang Yang, Lixin Zhou, and Nelly Panté. “Protein Myristoylation Plays a Role in the Nuclear Entry of the Parvovirus Minute Virus of Mice”. Journal of Virology 96 (2022), e01118–22.

[15] Rubén Sánchez-Eugenia, Julen Goikolea, David Gil-Cartón, Lissete Sánchez-Magraner, and Diego MA Guérin. “Triatoma virus recombinant VP4 protein induces membrane permeability through dynamic pores”. Journal of virology 89 (2015), 4645–4654.

[16] Rhinovirus VP4 Exhibit Cross-Serotypic. “Antibodies to the Buried N Terminus of”. J. Virol 83 (2009), 7040.

[17] Xiangxi Wang, Wei Peng, Jingshan Ren, Zhongyu Hu, Jiwei Xu, Zhiyong Lou, Xumei Li, Weidong Yin, Xinliang Shen, Claudine Porta, et al. “A sensor-adaptor mechanism for enterovirus uncoating from structures of EV71”. Nature structural & molecular biology 19 (2012), 424–429.

[18] Yue Liu, Ju Sheng, Arno LW van Vliet, Geeta Buda, Frank JM van Kuppeveld, and Michael G Rossmann. “Molecular basis for the acid-initiated uncoating of human enterovirus D68”. Proceedings of the National Academy of Sciences 115 (2018), E12209–E12217.

[19] Qingling Wang, Qian Yang, Congcong Liu, Guoqing Wang, Hao Song, Guijun Shang, Ruchao Peng, Xiao Qu, Sheng Liu, Yingzi Cui, et al. “Molecular basis of differential receptor usage for naturally occurring CD55-binding and-nonbinding coxsackievirus B3 strains”. Proceedings of the National Academy of Sciences 119 (2022), e2118590119.

[20] David Gil-Cantero, Carlos P Mata, Luis Valiente, Alicia Rodríguez-Huete, Alejandro Valbuena, Reidun Twarock, Peter G Stockley, Mauricio G Mateu, and José R Castón. “Cryo-EM of human rhinovirus reveals capsid-RNA duplex interactions that provide insights into virus assembly and genome uncoating”. Communications Biology 7 (2024), 1501.

[21] Chandra Shekhar Kumar, Debajit Dey, Sukanya Ghosh, and Manidipa Banerjee. “Breach: Host Membrane Penetration and Entry by Nonenveloped Viruses”. Trends in Microbiology 26 (2018), 525–537.

[22] Andrew Waterhouse, Martino Bertoni, Stefan Bienert, Gabriel Studer, Gerardo Tauriello, Rafal Gumienny, Florian T Heer, Tjaart A P de Beer, Christine Rempfer, Lorenza Bordoli, et al. “SWISS-MODEL: homology modelling of protein structures and complexes”. Nucleic acids research 46 (2018), W296–W303.

[23] Joshua D Yoder, Javier O Cifuente, Jieyan Pan, Jeffrey M Bergelson, and Susan Hafenstein. “The crystal structure of a coxsackievirus B3-RD variant and a refined 9-angstrom cryo-electron microscopy reconstruction of the virus complexed with decay-accelerating factor (DAF) provide a new footprint of DAF on the virus surface”. Journal of virology 86 (2012), 12571–12581.

[24] Hui Li, Andrew D Robertson, and Jan H Jensen. “Very fast empirical prediction and rationalization of protein pKa values”. Proteins: Structure, Function, and Bioinformatics 61 (2005), 704–721.

[25] Sunhwan Jo, Taehoon Kim, Vidyashankara G Iyer, and Wonpil Im. “CHARMM-GUI: a web-based graphical user interface for CHARMM”. Journal of computational chemistry 29 (2008), 1859–1865.

[26] Jing Huang, Sarah Rauscher, Grzegorz Nawrocki, Ting Ran, Michael Feig, Bert L De Groot, Helmut Grubmüller, and Alexander D MacKerell Jr. “CHARMM36m: an improved force field for folded and intrinsically disordered proteins”. Nature methods 14 (2017), 71–73.

[27] Jean-Paul Ryckaert, Giovanni Ciccotti, and Herman JC Berendsen. “Numerical integration of the cartesian equations of motion of a system with constraints: molecular dynamics of n-alkanes”. Journal of computational physics 23 (1977), 327–341.

[28] DA Case, HM Aktulga, K Belfon, IY Ben-Shalom, JT Berryman, SR Brozell, DS Cerutti, TEI Cheatham, GA Cisneros, VWD Cruzeiro, et al. “AMBER2020, university of California, San Fransisco”. J. Amer. Chem. Soc 142 (2020), 3823–3835.

[29] Paulo CT Souza, Riccardo Alessandri, Jonathan Barnoud, Sebastian Thallmair, Ignacio Faustino, Fabian Grünewald, Ilias Patmanidis, Haleh Abdizadeh, Bart MH Bruininks, Tsjerk A Wassenaar, et al. “Martini 3: a general purpose force field for coarse-grained molecular dynamics”. Nature methods 18 (2021), 382–388.

[30] Yoav Atsmon-Raz and D Peter Tieleman. “Parameterization of palmitoylated cysteine, farnesylated cysteine, geranylgeranylated cysteine, and myristoylated glycine for the martini force field”. The Journal of Physical Chemistry B 121 (2017), 11132–11143.

[31] Gary A. Huber and J. Andrew McCammon. “Browndye: A software package for Brownian dynamics”. Computer Physics Communications 181 (2010), 1896–1905.

[32] Huan-Xiang Zhou and Xiaodong Pang. “Electrostatic interactions in protein structure, folding, binding, and condensation”. Chemical reviews 118 (2018), 1691.

[33] Neeti Sinha and Sandra J Smith-Gill. “Electrostatics in protein binding and function”. Current Protein and Peptide Science 3 (2002), 601–614.

[34] Szilárd Páll, Artem Zhmurov, Paul Bauer, Mark Abraham, Magnus Lundborg, Alan Gray, Berk Hess, and Erik Lindahl. “Heterogeneous parallelization and acceleration of molecular dynamics simulations in GROMACS”. The Journal of chemical physics 153 (2020).

[35] Tsjerk A Wassenaar, Kristyna Pluhackova, Rainer A Bockmann, Siewert J Marrink, and D Peter Tieleman. “Going backward: a flexible geometric approach to reverse transformation from coarse grained to atomistic models”. Journal of chemical theory and computation 10 (2014), 676–690.

[36] Szilárd Páll, Mark James Abraham, Carsten Kutzner, Berk Hess, and Erik Lindahl. “Tackling exascale software challenges in molecular dynamics simulations with GROMACS”. International conference on exascale applications and software. Springer. 2014, 3–27.

[37] Naveen Michaud-Agrawal, Elizabeth J Denning, Thomas B Woolf, and Oliver Beckstein. “MDAnalysis: a toolkit for the analysis of molecular dynamics simulations”. Journal of computational chemistry 32 (2011), 2319–2327.

[38] Alan Grossfield. “WHAM: the weighted histogram analysis method, version 2.0. 9”. Available at membrane.urmc.rochester.edu/content/wham. Accessed November 15 (2013), 2013.

[39] Gareth A Tribello, Massimiliano Bonomi, Davide Branduardi, Carlo Camilloni, and Giovanni Bussi. “PLUMED 2: New feathers for an old bird”. Computer physics communications 185 (2014), 604–613.

[40] Rahul K Das and Rohit V Pappu. “Conformations of intrinsically disordered proteins are influenced by linear sequence distributions of oppositely charged residues”. Proceedings of the National Academy of Sciences 110 (2013), 13392–13397.

[41] Wolfgang Kabsch and Christian Sander. “Dictionary of protein secondary structure: pattern recognition of hydrogen-bonded and geometrical features”. Biopolymers: Original Research on Biomolecules 22 (1983), 2577–2637.

[42] Dale DO Martin, Erwan Beauchamp, and Luc G Berthiaume. “Post-translational myristoylation: Fat matters in cellular life and death”. Biochimie 93 (2011), 18–31.

[43] Anusha Panjwani, Amin S. Asfor, and Tobias J. Tuthill. “The conserved N-terminus of human rhinovirus capsid protein VP4 contains membrane pore-forming activity and is a target for neutralizing antibodies”. Journal of General Virology 97 (2016), 3238–3242.

[44] Jack WJ Welland, Henry G Barrow, Phillip J Stansfeld, and Janet E Deane. “Conformational dynamics and membrane insertion mechanism of B4GALNT1 in ganglioside synthesis”. Nature Communications 16 (2025), 5442.

[45] Juan Ortiz-Mateu, Diego Belda, Ana Isabel Avilés-Alía, Josep Alonso-Romero, Maria Jesús García-Murria, Ismael Mingarro, Ron Geller, and Luis Martinez-Gil. “The sequence and structural integrity of the SARS-CoV-2 Spike protein transmembrane domain is crucial for viral entry”. Communications Biology 8 (2025), 1579.

[46] Sergey V Ulianov, Artem K Velichko, Mikhail D Magnitov, Artem V Luzhin, Arkadiy K Golov, Natalia Ovsyannikova, Igor I Kireev, Alexey S Gavrikov, Alexander S Mishin, Azat K Garaev, et al. “Suppression of liquid–liquid phase separation by 1, 6-hexanediol partially compromises the 3D genome organization in living cells”. Nucleic acids research 49 (2021), 10524–10541.

[47] Ashutosh Shukla, Aditya K Padhi, James Gomes, and Manidipa Banerjee. “The VP4 peptide of hepatitis A virus ruptures membranes through formation of discrete pores”. en. J. Virol. 88 (2014), 12409–12421.

[48] Shantanu Guha, Jenisha Ghimire, Eric Wu, and William C. Wimley. “Mechanistic Landscape of Membrane-Permeabilizing Peptides”. Chemical Reviews 119 (2019). PMID: 30624911, 6040–6085.

